# Interictal activity fluctuations follow rather than precede seizures on multiple time scales in a mouse model of focal cortical dysplasia

**DOI:** 10.1101/2025.05.22.655485

**Authors:** Jan Kudlacek, Jan Chvojka, Michaela Kralikova, Salome Kylarova, Tejasvi Ravi, Ondrej Novak, Jakub Otahal, Martin Balastik, Premysl Jiruska

## Abstract

The unpredictability of seizure occurrence is a major debilitating factor for people with epilepsy. A seizure forecasting system would greatly improve their quality of life. Successful seizure forecasting necessitates a comprehensive understanding of the factors influencing seizure timing at multiple temporal scales. In this study, we investigated multiscale properties of interictal epileptiform discharges (IEDs) and seizure parameters in a highly realistic mouse model of focal cortical dysplasia-related epilepsy. We analyzed the properties’ evolution at four timescales, ranging from epilepsy progression and seizure clusters to circadian and peri-ictal changes. We discovered that the FCD-related epilepsy syndrome was progressive in terms of interictal activity rate and seizure characteristics. Sixty percent of seizures occurred in clusters. During the clusters, the seizure duration, seizure power, and IED rate were increasing. Circadian rhythm influenced seizure occurrence with the peak seizure probability at 4 p.m. under a standard 12/12 light dark cycle with lights-on at 6 a.m. Peri-ictal analysis revealed no significant change in IED rate preceding individual seizures; however, a consistent two-peak pattern of IED elevation was observed following seizures. Specifically, an initial peak in IED rate emerged 5–10 minutes post-seizure, returning to baseline within two hours, followed by a secondary peak 6–12 hours later, which again subsided to baseline levels in 24–48 hours. This pattern could be fitted with a sum of three exponentials. Using the three-exponential pattern, we simulated IED rate fluctuations in each animal. The smoothed simulated IED rates showed good agreement with the smoothed real recorded IED rates, suggesting that the cumulative effect of post-ictal IED patterns can account for long-term fluctuations in IED rate. Our results indicate that, in our model of FCD-related epilepsy, consistent IED rate fluctuations follow rather than precede individual seizures. Therefore, fluctuations in IED rate can be viewed as a reflection of cyclic seizure occurrence. This implies that either IED rate fluctuations or accurate seizure records may be equally valuable for seizure risk forecasting.

**Highlights:**

- FCD-related epilepsy model displays a progressive nature and fluctuations between high and low seizure risk.
- Seizures occur with higher probability in the day time which corresponds to sleep-related seizures commonly occurring in human patients with FCD.
- IED rate increases significantly after seizures, indicating a postictal effect rather than a preictal one, with the post-ictal phase displaying a two-peak pattern of fast and slow IED rate increase.
- Long-term changes in the IED rate could be attributed to the time-dependent cumulative effect of the two-peak seizure-related increase in IED rate.

## Introduction

Epilepsy is a chronic neurological disorder affecting 0.5 to 1 % of population (Beghi, 2020, Ngugi et al., 2010). It is characterized by spontaneous recurrent seizures which can be pharmacologically controlled in less than two thirds of patients (Chen et al., 2018). Importantly, the number of drug-resistant patients has not changed significantly with the introduction of new anti-seizure medications. Other therapies such as surgery, neurostimulation or ketogenic diet are only suitable to a small population of people with drug-resistant epilepsy and the results are often less than optimal (Fattorusso et al., 2021). Therefore, there is a need for novel tools for seizure management, including approaches for seizure prediction or forecasting which could decrease the fear from suddenly occurring seizures (Fisher et al., 2000) and help optimize therapies (Ramgopal et al., 2013). Although sophisticated mathematical analyses of intracranial EEG (iEEG) showed some promise for predicting individual seizures, seizure prediction has not yet been introduced to the clinical practice (Lehnertz et al., 2023). The seizures are, however, not completely random, i.e. they are not a Poisson process. This fact gives hope for another approach called seizure forecasting which aims at informing the patient about their seizure risk rather than individual approaching seizures (Baud and Rao, 2018; Karoly et al., 2020).

The non-Poissonian nature of seizure occurrence is evident on multiple levels. First, the seizure rate may be increasing (Gorter et al., 2001; Raedt et al., 2009; Williams et al., 2009) or decreasing (Behr et al., 2017) over lifetime. Second, seizure rate fluctuates in multiday cycles in most patients (Balish et al., 1991; Bauer and Burr, 2001; Griffiths and Fox, 1938; Sunderam et al., 2007; Tauboll et al., 1991). This has been confirmed using continuous chronic intracranial EEG (iEEG) recordings spanning durations of months to years (Baud et al., 2018; Karoly et al., 2021; Maturana et al., 2020). The fluctuations can lead to the formation of seizure clusters interspersed by periods of low seizure risk or even seizure freedom. Importantly, this slow dynamics of seizure occurrence and clustering observed in patients is replicated in various animal models of epilepsy, including models of temporal lobe epilepsy (TLE) in rats (Bajorat et al., 2011; Dudek and Staley, 2011; Eskikand et al., 2025; Goffin et al., 2007; Grabenstatter et al., 2005; Hawkins and Mellanby, 1987; Kudlacek et al., 2021) and mice (Mazzuferi et al., 2012), neocortical epilepsy induced by tetanus toxin (B. L. Chang et al., 2018) or by perinatal hypoxia (Kadam et al., 2010), and in spontaneously epileptic dogs (Gregg et al., 2020). Fluctuations of seizure rate were also apparent in a murine model of focal cortical dysplasia (Almacellas Barbanoj et al., 2024) but were not investigated in detail. In a model of TLE, seizure duration was also reported to linked to cycles of seizure rate (Eskikand et al., 2024).

Interictal epileptiform discharges (IEDs) are a hallmark of epileptic tissue. IED rate is known to fluctuate in circadian and multidien cycles in both human patients (Leguia et al., 2021; Maturana et al., 2020) and animal models of epilepsy (Baud et al., 2019; Kadam et al., 2010; Levesque et al., 2011). Seizures were shown to occur with the highest probability on the rising phase of the IED rate cycles giving rise to forecasting algorithms based on non-causal estimation of IED rate cycle phase (Proix et al., 2021). The multidien cycle period, however, varies from cycle to cycle (Schroeder et al., 2023) which complicates the causal estimation of the cycle phase. This limitation was recently overcome by estimating the phase of the cycle causally using hippocampal functional connectivity patterns in a cohort of patients with bitemporal epilepsy. This approach enabled pseudo-prospective forecasting of seizure risk for the first time (Khambhati et al., 2024). Similar studies on neocortical epilepsy are, however, missing.

Circadian distribution of seizures has been well studied in patients and animal models of epilepsy. Analysis of continuous long-term iEEG recordings from people with intractable epilepsy confirmed that seizures and IEDs follow a circadian patterns with the phases if IED rate and seizure rate circadian fluctuations often aligned (Karoly et al., 2016). Moreover, neocortical seizures tended to peak during the night whereas temporal lobe seizures during the day (Spencer et al., 2016) which is in line with the data from presurgical intracranial EEG monitoring which showed that probability of seizures originating in frontal and parietal cortices peaked between 4 a.m. and 7 a.m. (Durazzo et al., 2008) whereas temporal lobe seizures were most likely to occur in the afternoon. In status epilepticus-based models of TLE, some studies reported no circadian variation of seizure likelihood (Bajorat et al., 2011) and in others, seizures were more common during the light period (Baud et al., 2019; Bertram and Cornett, 1994; Gorter et al., 2001; Hellier and Dudek, 1999; Pitsch et al., 2017; Quigg et al., 1998; Raedt et al., 2009) which corresponded to lower activity phase of the day since mice and rats are nocturnal animals. In the intrahippocampal tetanus toxin model of TLE, no circadian pattern of seizure occurrence was observed (Eskikand et al., 2025) and dogs with spontaneously occurring epilepsy presented subject-specific times of peak seizure occurrence (Gregg et al., 2020). In a model of FCD, seizures were more prevalent during the day (Almacellas Barbanoj et al., 2024).

A possible seizure prediction strategy seeks for changes in IED rate ahead of seizures. Although this strategy has so far never led to successful seizure prediction, patient-specific increases or decreases of IED rate in the preictal period were occasionally reported in patients (Karoly et al., 2016). No such data exist from a murine model of FCD.

In this study, we explored the multiscale temporal distribution of seizures and accompanying changes in IED rate to disclose the underlying disease dynamics in a murine model of FCD-related neocortical epilepsy. We identified that seizures induced early (minutes) and delayed (hours) increases in IED rate. The cumulative effect of seizure-related IED facilitation replicates the long-term IED profile and allows estimation of the future evolution of IED rate.

## Methods

### Animals

We analyzed long-term EEG recordings from 13 C57BL6/N mice (9 males and 4 females) with experimentally induced focal cortical dysplasia (FCD). The mice were housed under standard conditions in a room with controlled temperature (22±1°C) and 12/12 h light/dark cycle and had *ad libitum* access to food and water. All animal experiments were performed under the Animal Care and Animal Protection Law of the Czech Republic, fully compatible with the guidelines of the European Union directive 2010/63/EU.

### FCD induction

The FCD lesion was induced by *in utero* electroporation (Saito and Nakatsuji, 2001; Tabata and Nakajima, 2001). Pregnant mice (day 14.5 ± 0.5 post-fertilization) were anesthetized with isoflurane. Uterine horns were exposed by midline laparotomy, and the right lateral ventricle of each embryo was injected with a mixture of plasmids diluted in 2 μg/ml Fast Green (F7252, Sigma, USA), using pulled and beveled glass capillaries. The plasmid mixture contained two plasmids: 3 μg/μl of pCAG-mTOR-IRES-mAmetrine where the mTOR had the mutation p.Leu2427Pro to induce the FCD (SoVarGen Co., Ltd., Korea; Lim et al., 2015) and 1.5 μg/μl of pCAG-EGFP to visualize the lesion (Figure 1A-C). After injection, forceps-type electrodes (3 mm, CUY650P3, Nepa Gene) were positioned on the head of each embryo, and electroporation of plasmids was elicited by NEPA21 (Nepa Gene, Japan) electroporator delivering five 35 V, 50 ms pulses, with 950 ms interpulse intervals (Procházková et al., 2024).

**Figure 1:**
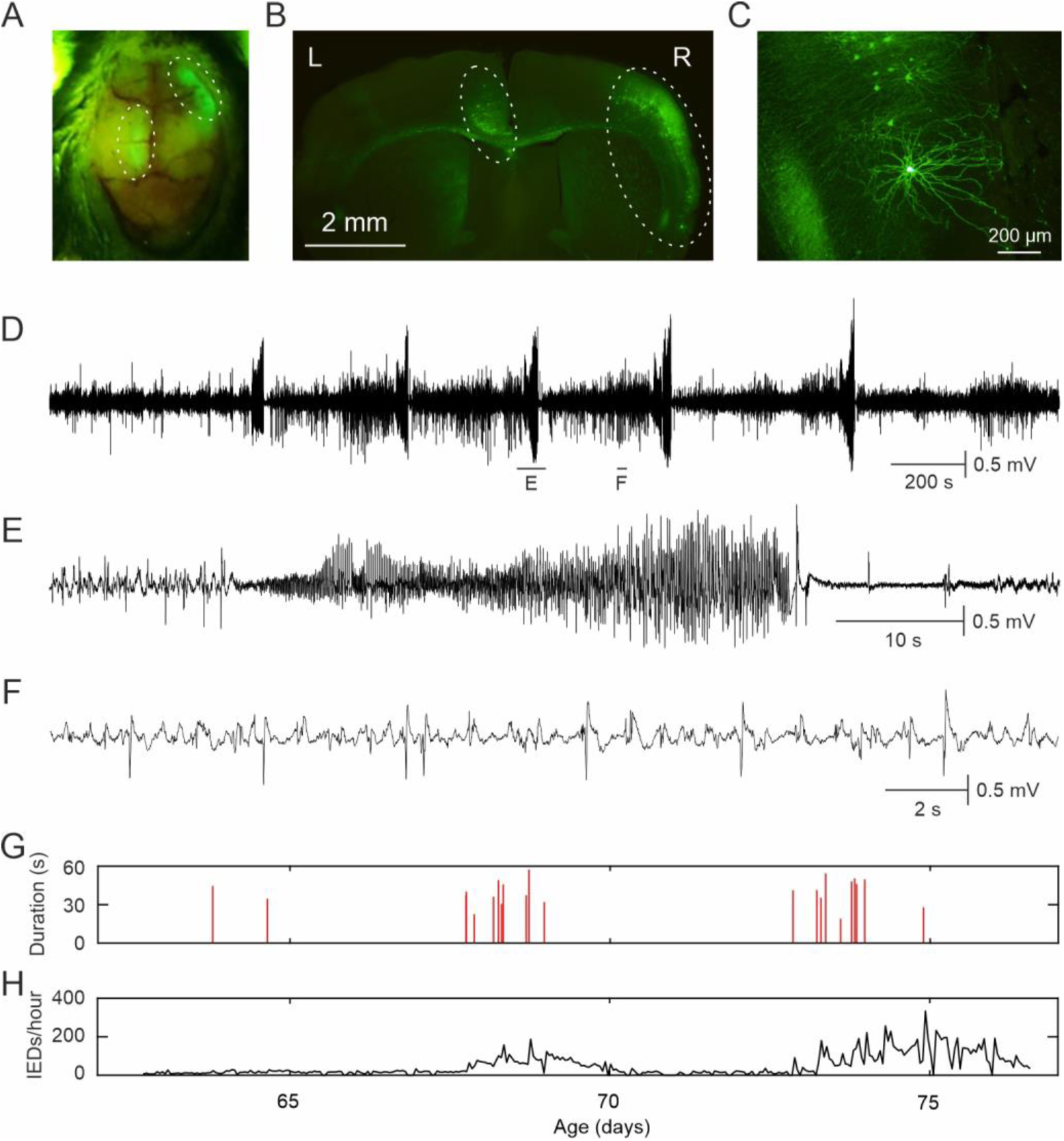
Basic features of the mouse model of FCD. (A) View of the mouse skull before the surgery with the GFP fluorescence-marked FCD lesion in the right frontal cortex. (B) Coronal slice from the same mouse reveals two areas of dysplastic cortex with GFP-positive cells scattered across the cortex and the white matter. (C) An example of a dysmorphic neuron characterized by large body and abnormal dendritic branching. (D) EEG recording of a cluster of five seizures. (E) Detailed example of a seizure and interictal epileptiform discharges (F). (G) Seizure profile showing times of occurrence and durations of seizures. (H) Corresponding IED rate profile.

### Electrode implantation and EEG recording

Mice were implanted with EEG electrodes at 6–10 weeks of age. Prior to implantation, custom EEG implants were prepared by soldering coated silver wires (127 μm in diameter, AM Systems, Inc., USA, cat. no. 786000) to prefabricated connectors (TME Electronic Components, Poland, cat. no. DS1065-03-2*6S8BV). Mice were anesthetized with isoflurane, a small area of skin on the head was removed to expose the frontal and parietal bones of the skull, and GFP fluorescence macroscopic imaging was performed to determine the position of the FCD (Figure 1A). Five holes were drilled through the skull. Four epidural electrodes were implanted bilaterally in the frontal and parietal bones at approximate coordinates from bregma AP: ±1.5 mm, L: ±1.5mm. One hole was drilled above the cerebellum for grounding/reference electrode. The electrodes together with the connector were attached to the skull using cyanoacrylate glue (Loctite, USA, cat. no. 1363589). Following a five-day recovery period, the animals were individually video-EEG monitored continuously for >2 weeks using custom-made hardware and software. The signal was amplified directly on the mouse head using a headstage amplifier, filtered between 1 and 500 Hz and digitized at 2 kHz sample rate.

### EEG analysis

EEG was analyzed using custom-written scripts in Matlab 2024b computing environment (Mathworks Inc., USA). First, data were downsampled to 250 Hz sample rate. Seizures (Figure 1D, E) were identified by manual review of the EEG. Interictal epileptiform discharges (IEDs; Figure 1F) were detected using a published detector (Janca et al., 2014) with the default settings. IEDs from the same or different channels occurring less than 100 ms apart were considered one event. The EEG was contaminated by electromyographic (EMG) artifacts, which were occasionally mistaken for IEDs by the detector. Therefore, the EMG-contaminated epochs were excluded from the analysis of IED rate (see below).

### Electromyographic artifact detection

The electromyographic (EMG) artifacts were automatically detected by a custom-made detector based on the following algorithm. For the EMG detection, we used the original data sampled at 2 kHz. For each mouse, four 5-minute EEG files were randomly selected (two from the daytime period and two from the nighttime period) and the EMG artifacts were manually labeled. Then, signal features of the EMG-contaminated and clean epochs were computed in 0.5 second bins. The features were signal variance, entropy, sum of amplitude spectrum in four bands (2 – 10 Hz, 10 – 50 Hz, 50 – 200 Hz, 200 – 800 Hz), entropy of phase spectrum and sum of phase spectrum difference which aimed to make the detector sensitive not only to the frequency content but also to shape of the signal, i.e. random noise (EMG) vs. high-amplitude spike (IED). Multiple classifiers (Coarse Tree, Medium Tree, Boosted Trees and Bagged Trees) were then trained using Matlab’s built-in Classification Learner application or built-in functions. For each mouse, we selected the one that yielded the highest accuracy in the cross-validation procedure. In the whole recording, the signal features were then computed and the classifier applied to automatically label the EMG-contaminated epochs.

### Analysis of trends in seizure characteristics

For the seizures, we analyzed their rate of occurrence, duration and mean EEG signal power (see example in Figure 1G). In order to evaluate a possible trend in the characteristic, we divided the analyzed period into bins. In case of the analysis of the progression over the whole recording period, the bins were 24-hour long with edges at midnight. The first and last bins were shorter since the recording never commenced or concluded at midnight. In case of a signal dropout, the length of the bin is equal to the total time of the valid signal only. Then, we computed the mean of the characteristic in the given bin. In the analysis of the seizure rate, we took into account the length of the bin. The bin means were then fitted with a straight line using the least squares method (Matlab function `fit`). For the fitting of characteristics other than seizure rate, the bins were weighted by their length so that the shorter bins had proportionately lower influence on the fit.

In the interictal signal, IED rate was analyzed in 1-hour or 1-minute bins, and trends were analyzed similarly. For each bin, we accounted for the actual duration of usable signal, which was often shorter than the bin length due to signal dropouts or contamination from electromyographic (EMG) artifacts.

### Statistical evaluation of trends

The statistical evaluation of increasing or decreasing trends in each characteristic was performed by the following procedure. We fitted a straight line to the binned data of each period of interest (whole recording or periods related to seizure clusters or seizures, see Results). The slopes of the lines were extracted. If multiple periods of interest occurred in a subject (e.g. multiple clusters), the slopes within each subject were averaged, thus giving one number per subject. Subject-wise statistics was then calculated using the Wilcoxon signed-rank test to determine whether the slopes were statistically significantly different from zero. Since we analyzed four characteristics, the p-values were subjected to Benjamini-Hochberg false discovery rate correction at the false discovery rate 0.1. The circadian rhythm-related analyses were computed using MATLAB Circular Statistics Toolbox (Berens, 2009).

### IED rate simulation

The IED rate after the seizures showed a consistent two-peak shape which could be modelled by a sum of exponentials. At the shorter time scale of two hours, we fitted the after-seizure IED rate by a sum of two exponentials (expression ‘A1*exp(k1*x)+A2*exp(k2*x)’ with the independent variable x). At the longer time scale of two days, we used three exponentials (’A1*exp(k1*x)+A2*exp(k2*x)+A3*exp(k3*x)’). The fitting was performed by the least squares method using Matlab function fit. Then, we leveraged the knowledge of the shape of IED rate after seizure to simulate artificial IED rate based solely on the seizure occurrence times. For this purpose, we designed an IIR filter with the impulse response equal to the fitted exponentials. Thus, the impulse response of the filter is

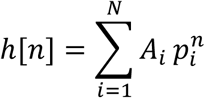

where *n* is discrete time index, *i* is index of fitted exponential, *N* is number of exponentials *A*_*i*_ is amplitude of *i*^*th*^ exponential and *p*_*i*_ is the *i*^*th*^ pole of the filter which is computed from the fitted values of *k*_*i*_ and sample period *T*_*s*_ as

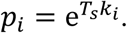

The filter transfer function in the z-transform is then

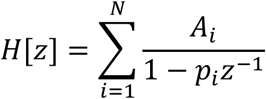

This can be expanded to

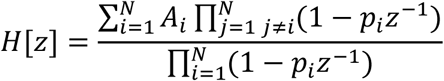

By expanding the numerator and denominator we convert the expression to the form of

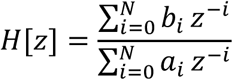

where *b*_*i*_ and *a*_*i*_ are the feedforward and feedback coefficients of the filter, respectively.

The obtained filter was then applied on the seizure rate evaluated in the same time bins as the IED rate (1-minute). The seizure rate signal contained zeros most of the time with occasional ones and rarely higher values marking the seizure occurrence. The result of the filtration was regarded as the simulated IED rate. Both original and simulated IED rates were smoothed by a 4-hour moving average. The similarity of the two IED rates was then evaluated using Pearson correlation coefficient and Relative error, which is essentially a normalized root mean squared error computed according to the formula

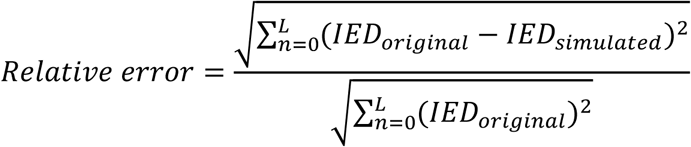

## Results

### Seizure profiles

The data are presented as means and medians in form mean±SEM (median±interquartile range). We analyzed data from 14 mice with FCD-related epilepsy, which developed seizures and interictal activity (Figure 1). The mice were monitored for 37±7 (32±44) days. During this period, individual mice experienced 87±30 (42±54) seizures and the mean seizure frequency was 2.4±0.4 (1.9±0.8) seizures/day. Mean seizure duration was 51±3 (48±15) s. Visual inspection of the seizure profiles in a raster plot (Figure 2) revealed fluctuations of seizure rate and seizure clusters. Since the mean seizure rate differed between the mice and the cluster durations were variable we used a statistical, data-driven definition of a seizure cluster. We defined seizure cluster as a group of at least four seizures separated from other seizures by at least 2⋅ISI_cluster max_, where ISI_cluster max_ stands for the longest inter-seizure interval within the cluster. Apart from that, the cluster had to be separated > 2⋅ISI_cluster max_ from the beginning and end of the recording. The cluster was also removed from analysis if it contained a significant signal dropout or was separated < 2⋅ISI_cluster max_ from a significant signal dropout. A signal dropout was considered significant if it was longer than both the IED analysis window (1 hour) and the shortest inter-seizure interval of a given mouse. This definition allows for the existence of nested clusters. We observed nesting over up to four levels (Figure 2, inset). We identified 12±5 (5±7) clusters per mouse. When only non-nested clusters were taken into account it was 6.7±3.3 (2.5±2) clusters per mouse. In the subsequent analyses, the nested clusters were included unless stated otherwise. The clusters lasted 18±3 (17±15) hours and contained 8.0±0.8 (7.4±3.4) seizures. Out of all seizures, 59±6 (59±43) % occurred within a cluster. The between-cluster period was 6.9±3.0 (3.5±2.9) days (only non-nested clusters considered). Note that during the between-cluster period, there could have occurred non-clustered seizures.

**Figure 2:**
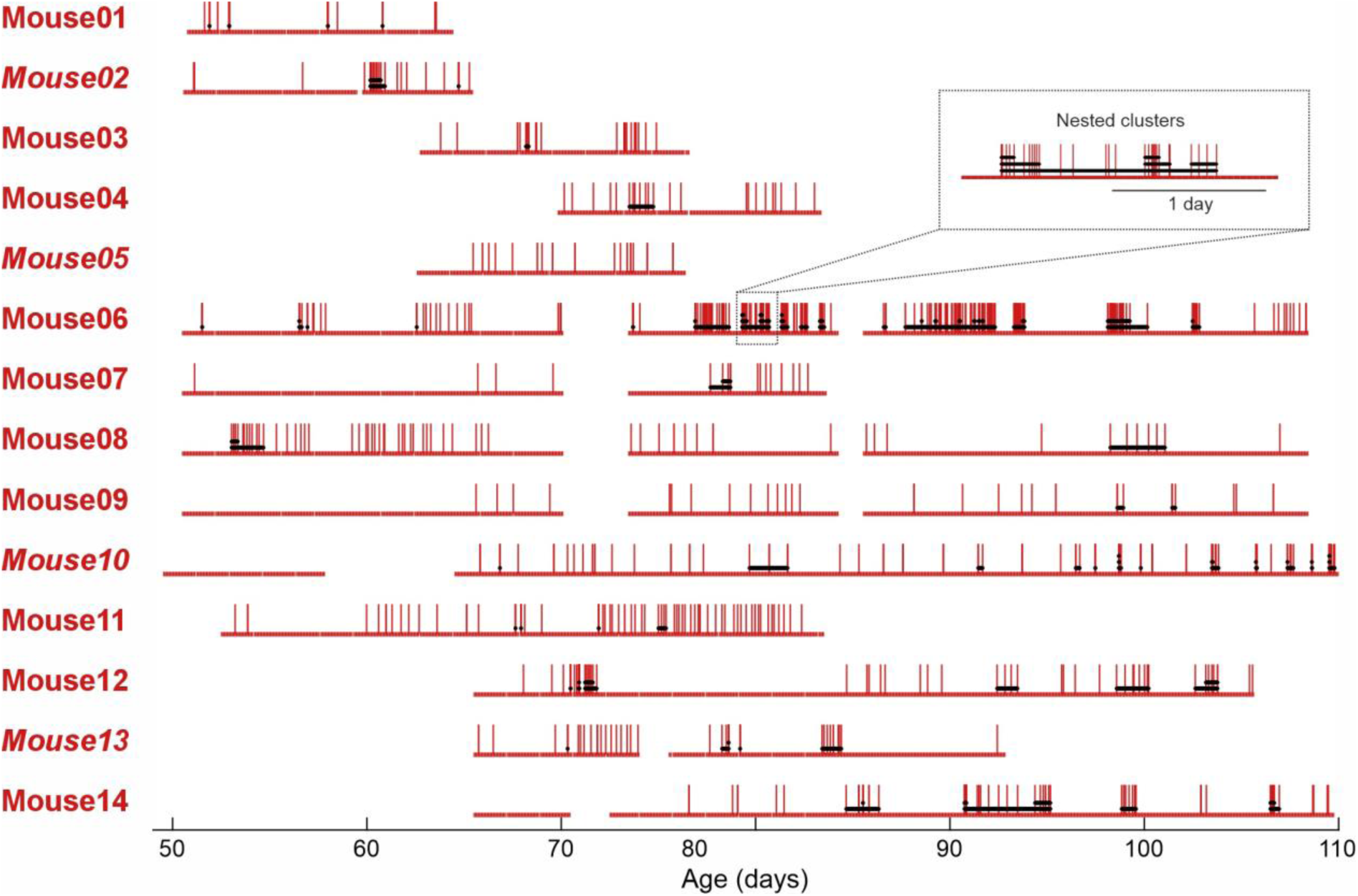
Seizure profiles. Red vertical lines indicate seizures. Red horizontal line at the bottom of each animal’s row indicates the presence of recording; gaps indicate recording dropouts due to technical failures. Black horizontal lines mark seizure clusters. Note the cluster nesting, sometimes over multiple levels (example in the inset). Female mice are marked by italic font.

### Life-long progression of the epileptic phenotype

First, we evaluated whether the FCD-related epilepsy in our model has a progressive nature. We divided the whole recording of each mouse into 24-hour time bins from midnight to midnight, except for the first bin which was from the first recorded seizure to the first midnight and the last bin which was from the last midnight to the last recorded seizure (Figure 3A). We calculated seizure rate, mean seizure duration and mean seizure signal power in these bins. IED rate was calculated in 1-hour long bins (Figure 3B). The trends were evaluated by fitting a line to the bin values and processed according to the Methods.

**Figure 3:**
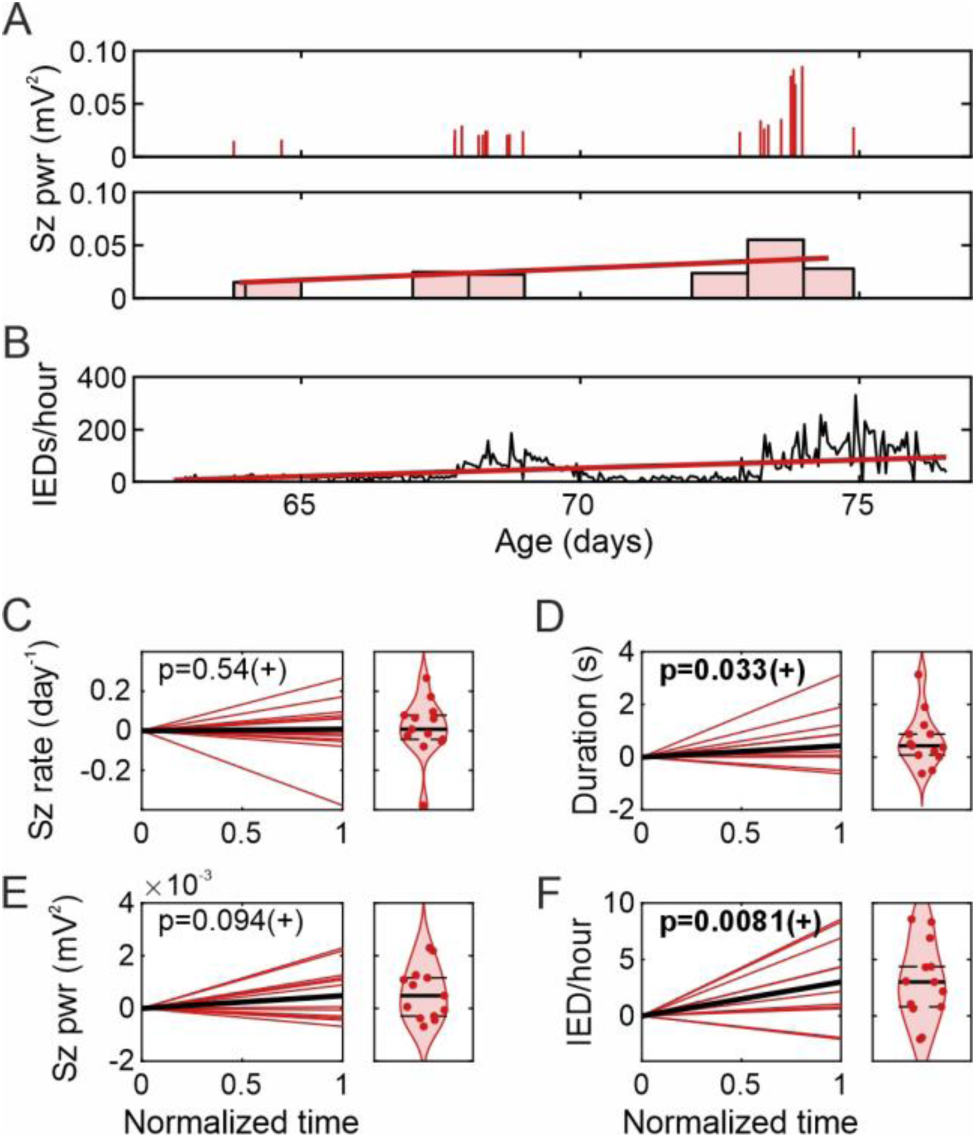
Long-term progression of epileptic phenotype. (A) Top: An example characteristic of seizures, the seizure signal power, is shown as the height of the lines. Bottom: The characteristics were averaged in 1-day-long bins from midnight to midnight (bar graph) and the bin values were fitted with a straight line (red solid line). Note that the first and last bins are shorter since the analyzed period was defined by the first and last recorded seizure and these never happened exactly at midnight. (B) The IED rate was analyzed in 1-hour bins (black line) and fitted with a straight line (red line). (C-F) Slopes of the fitted lines of individual animals (red) and median slope (black) with corresponding violin plots. The units of the slopes are the change per day of the quantity indicated to the left (e.g. the change of IED rate (IEDs/hour) per day). The y-axis in the violin plots is the same as in the line plots. The solid line in the violin plot marks the median, and dashed lines mark the bootstrapped 95% confidence interval. P-values of the Wilcoxon signed rank test are shown. The sign in the parentheses indicates if the median trend was positive or negative. Statistically significant p-values after FDR correction are in bold.

The plots for all individual mice are in Supplementary Figure 1. The seizure rate showed occasional increases or decreases in 4/14 individual animals but the population data do not show any consistent trend (p=0.54, Figure 3C). The seizure duration was, however increasing in 8/14 animals and also the population data show a significant increase (p=0.033, Figure 3D). The signal power shows only mild indication of an increase (p=0.094, Figure 3E). The most prominent progression was found in the IED rate was increasing in 12/14 animals and in population data (p=0.0081, Figure 3F). Thus, the syndrome appeared progressive in terms of seizure duration and interictal activity but not seizure rate. In none of the mice we observed indications of seizure remission.

### Seizure clusters

Next, we analyzed the same characteristics on a shorter timescale defined by the naturally occurring clusters of seizures. The seizure characteristics were binned in four equal bins per cluster (Figure 4A) and the bin values were fitted with a straight line (Figure 4B). Then, the slopes of the lines were averaged over clusters within each animal, and then, population statistics was computed (Figure 4C-E). The IED rate in 1-hour bins was fitted not only during the cluster but also in the pre-cluster and post-cluster periods of the same duration as the cluster (Figure 4F-H). Note, that nested clusters were included in this analysis. Similarly to lifelong progression, the data do not show any consistent trend in the seizure rate (p=1, Figure 4C). Along the seizure clusters, we observed an increase of seizure duration (p=0.021, Figure 4D), seizure signal power (p=0.027, Figure 4E), and IED rate (p=0.0034, Figure 4F). No significant change in IED rate was observed ahead of clusters (p=0.74, Figure 4G) and an indication of a decrease was observed after the clusters (p=0.08, Figure 4H). The data for individual mice are in Supplementary Figure 2. Apart from that, we compared the duration and signal power of the cluster terminating seizures (last seizure of a given cluster) to the other seizures within the respective clusters. The non-terminating seizures had the duration of 52±3.8 (53±16) seconds whereas the terminating seizures had the duration of 57±4.8 (58±23) (p=0.027, Wilcoxon signed rank test). The seizure signal power of the non-terminating and terminating seizures was 0.039±0.0048 (0.041±0.021) and 0.042±0.0057 (0.043±0.029), respectively, which was not significantly different (p=0.018, Wilcoxon signed rank test).

**Figure 4:**
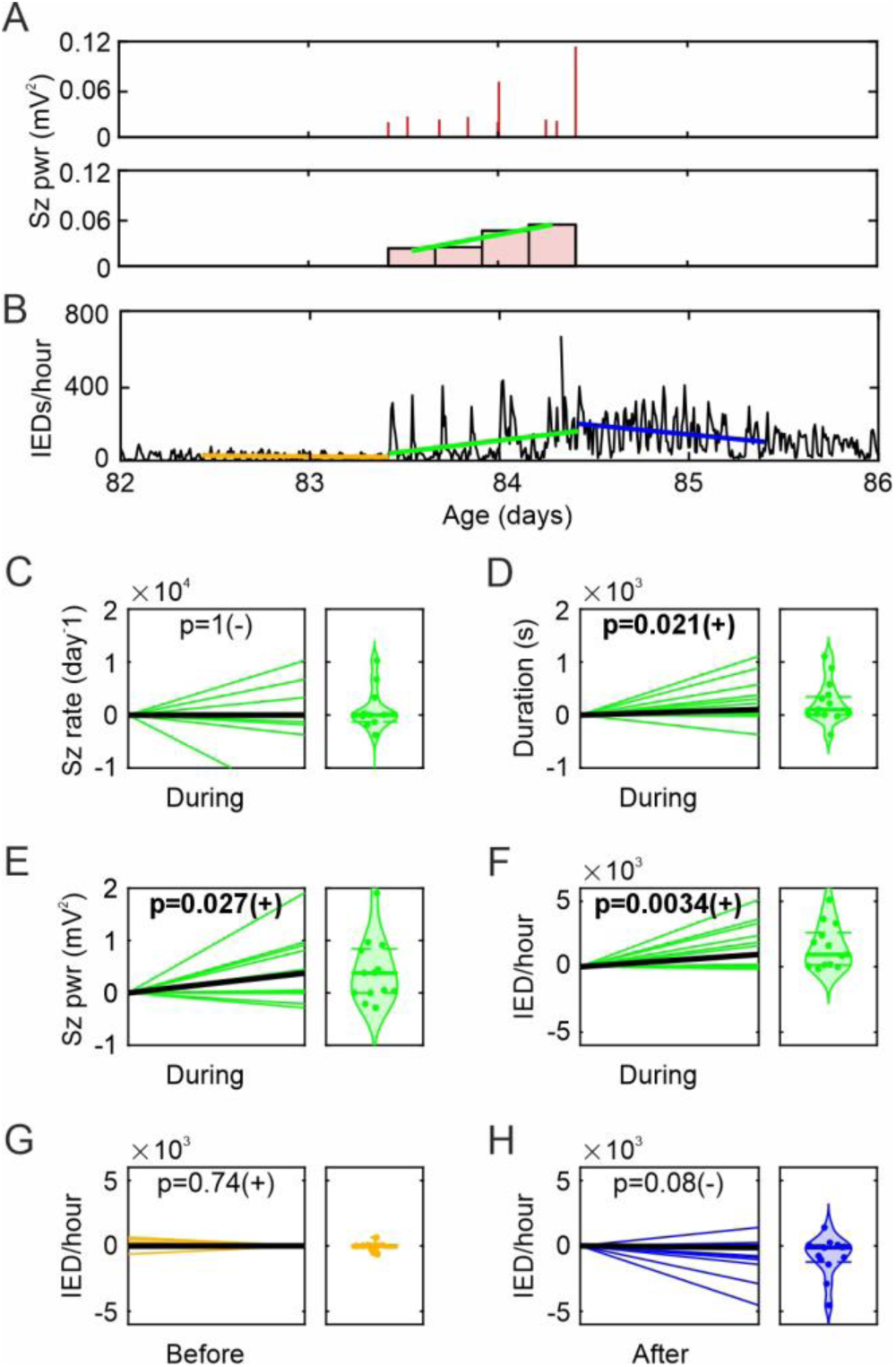
Analysis of trends before, during and after seizure clusters, color-coded by orange, green and blue, respectively. (A) An example of the analysis of a seizure cluster from Mouse 13. Seizure data were analyzed similarly to Figure 3 but with a different bin duration, in this analysis always four equal bins per cluster. (B) IED rate was analyzed in 1-hour bins, same as in Figure 3. In this example there is no change before, an increase during and a decrease after the cluster. The pre- and post-cluster periods are defined to have the same duration as the cluster. For each mouse, we averaged the slopes from all of its clusters. Thus, for each characteristic, we got one value per mouse. (C-G): Mean slopes from individual mice (color) and median slopes across subjects (black). The units are the change of the indicated quantity per day. P-values of the Wilcoxon signed rank test are shown. The sign in the parentheses indicates if the median trend was positive or negative. Statistically significant p-values after FDR correction are in bold.

### Circadian rhythm

The next timescale, on which we analyzed the data, was the circadian one. For the analysis of the seizure characteristics, we divided the day into 8 time bins. The seizure data were then binned according to the time of the seizure’s occurrence while ignoring the date (Figure 5A, B). The IED rate was averaged similarly in 24 bins around the day (Figure 5E). From the binned data of each mouse, we calculated the mean resultant vector (Figure 5A, E), i.e. the vector sum of all bin values while considering the angle of the bin center and bin value as the modulus. Then, from the set of resultant vectors, we calculated the population resultant vector which pointed at at 4 p.m. with the radius r=0.6 (Figure 5C). We applied the Rayleigh test of circular distribution non-uniformity on the angles of the individual resultant vectors. The results showed that most animals had the highest seizure likelihood in the afternoon and early night (p=0.008, Rayleigh test, Figure 5C, D). Surprisingly, we did not observe any consistent circadian modulation of the IED rate (Figure 5F, G) although in 10/14 individual animals, the circadian modulation was statistically significant (Supplementary Figure 3). We also tested if the seizure duration and signal power are modulated by the circadian rhythm but we did not find any trend (Supplementary Figure 3).

**Figure 5:**
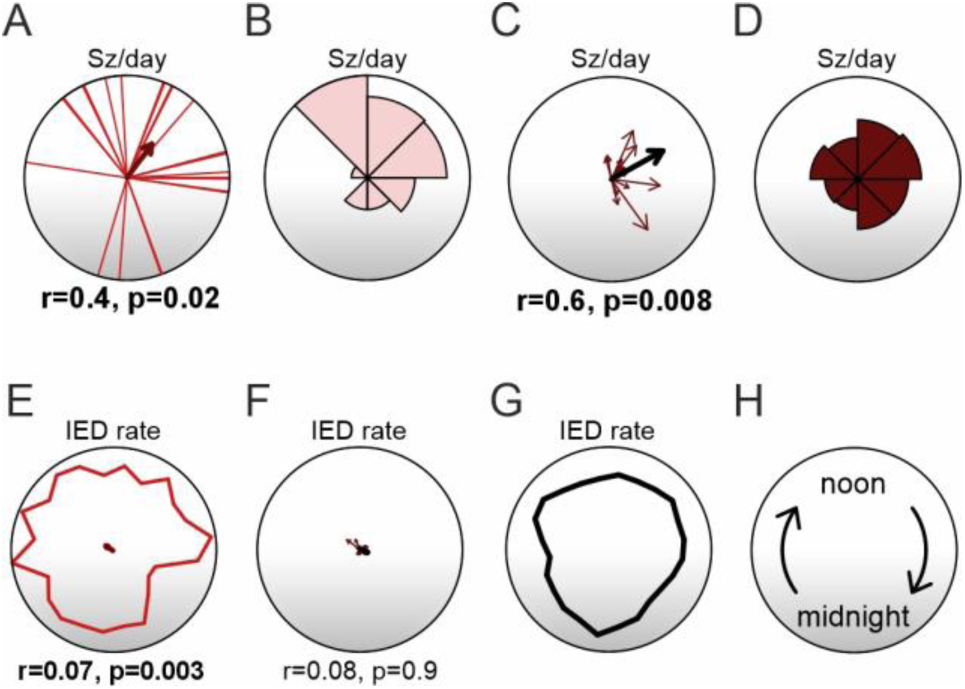
Circadian variation of seizure characteristics and IED rate. In each circular graph, midnight is at the bottom, noon at the top and time goes clockwise (H). (A) Example of seizure distribution from Mouse 5. Each red line marks the time of day of a seizure (regardless of date). The dark red arrow shows mean resultant vector (the radius of the circle is one). Below the graph, the length of the mean resultant vector r and the p-value of the Rayleigh test are shown. (B) Normalized circular histogram of seizure rate of Mouse 5. (C) Mean resultant vectors of all mice (dark red) and the population mean resultant vector created from those vectors (black). Radius of the circle is one. Below is the length of the mean resultant vector r and the p-value of the Rayleigh test. Note that for the Rayleigh test, only the directions of the mice’s mean resultant vectors were used. (D) Average of normalized circular histograms of all mice. (E) Circadian distribution of IED rate (red) with mean resultant vector (dark red) with its length r indicated below and p-value of Rayleigh test for Mouse 5. (F) Mean resultant vectors of IED rates of all mice and population mean resultant vector with its length r and Reyleigh test p-value below. Note that the mean resultant vectors are very short due to almost uniform IED rate distribution. (G) Mean circadian distribution of IED rate over all mice. (H) Explanatory figure for the time orientation. The lights were switched on at 6 a.m. and off at 6 p.m.

### Peri-seizure changes in IED rate

On the shortest timescale, we analyzed pre-seizure and post-seizure IED rate curves (Figure 6). Here, the IED rate was analyzed in 1-minute time bins, to capture more detail in the temporal profile of IED rate. Since, it was not obvious how fast the pre- and post-ictal changes can be expected we analyzed two durations of before- and after-seizure periods: 2 hours and 2 days.

**Figure 6:**
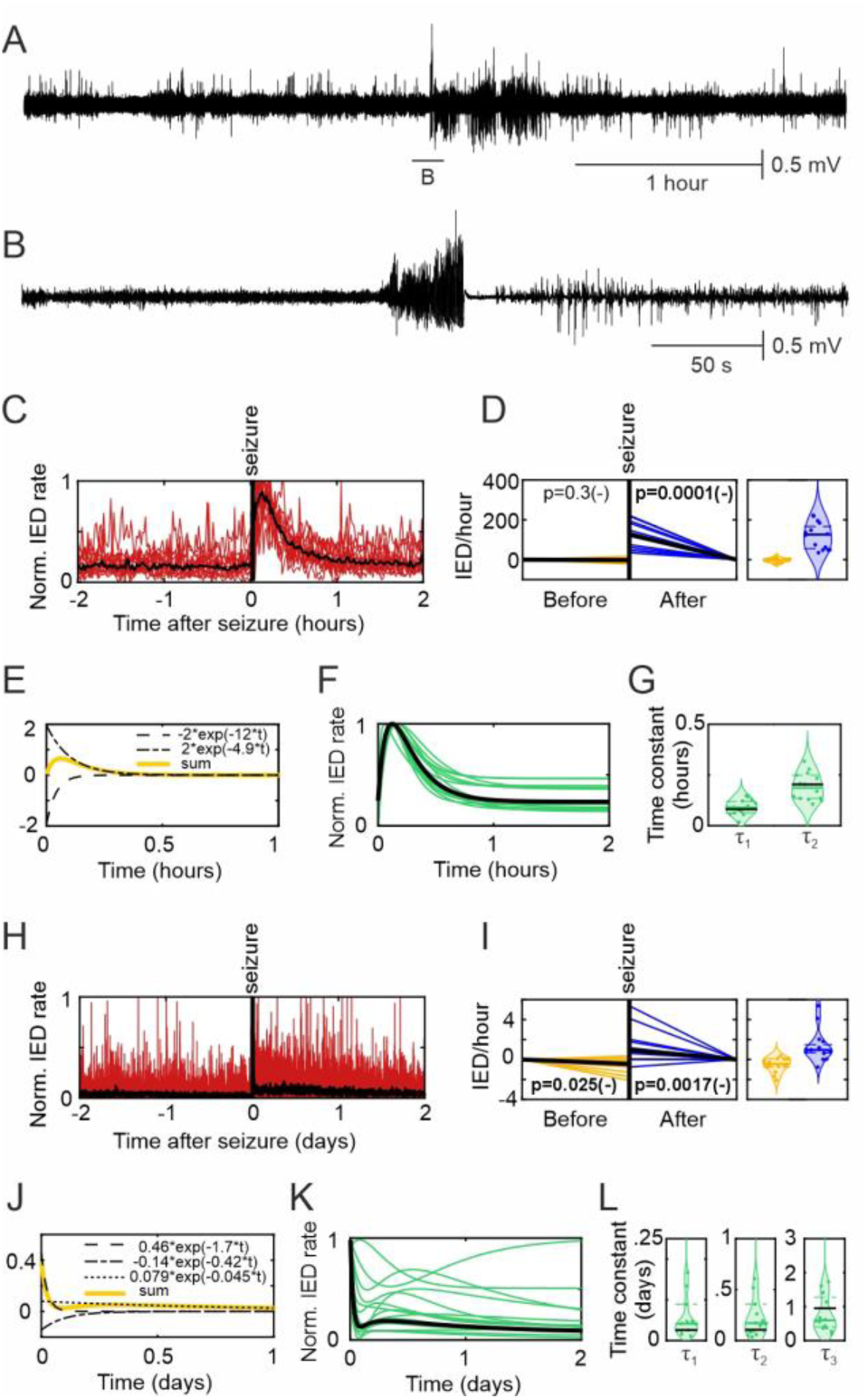
IED rate trends before and after seizure. (A) Seizure with two hours before and two hours after it. Increased interictal activity is visible after the seizure. (A) Detail of the seizure from A with three minutes before and after. (C) IED rate before and after a seizure was analyzed in 1-minute bins, averaged across all seizures within each mouse and normalized to the maximum value within the mouse (red lines). Median across the mice is shown by the black line. (D) Straight lines were fitted to two-hour segments before and after each seizure. Mean slopes over all seizures of each mouse are shown in yellow (before seizure) and blue (after seizure). The units are the change of IED rate in IEDs/hour per day. Note that after the seizure, negatives of the slopes are plotted at the seizure offset time and zero at the end of the fit because it corresponds visually to the raw data in (C). Correspondingly, the blue violin plot shows negatives of the slopes. The y-axis for the violin plots is the same as for the line plots. (E) The sum of two exponentials was fitted to the after-seizure IED rate. (F) Green curves: Fits for each mouse’s post-seizure IED rate (i.e. fits to the red curves in C). Black curve: Fit to the median post-seizure IED rate over all mice (i.e. to the black curve in C). (G) Violin plots indicating the time constants of the two exponentials for each mouse. The solid green horizontal line indicates the median and the green dashed lines the bootstrapped 95% confidence intervals of the individual mice data. The black horizontal line indicates the values from the fit of the averaged data (black line in F). (H-L) Analyses for the longer fitted segment (two days here) and fitted curve consisting of the sum of three exponentials.

In the case of the 2-hour period before the seizure, we required the separation of the analyzed seizure from the previous one to be at least four hours. Similarly, the after-seizure period was only included if the next seizure occurred at least 4 hours after the analyzed one. A similar safety margin was, however, not applied in the 2-day period analysis since very few mice showed seizure-free periods longer than 2 days. Therefore, in this case we only required the analyzed 2-day-long before- or after-seizure period to be seizure-free.

Most postictal periods displayed an increase in the IED rate in the raw signal (Figure 6A, B). In the IED rate curve, a transient fast increase in IED rate was observed approximately 10 minutes after the seizure in both individual mice data (Supplementary Figure 4) and averaged population data (Figure 6C). No change in the IED rate was observed before the seizure. When the IED rate curves were fitted with straight lines, their slopes were not different from zero before (p=0.3), but clearly negative after the seizure (p=0.0001, Figure 6D). The 2-hour after-seizure curve could be fitted with a sum of two exponentials (Figure 6E) both in individual mice (Supplementary Figure 5) and averaged data (Figure 6F). The time constants for the rising exponentials were τ_1_ = 5.4 ± 0.6 (5.4 ± 3.5) and for the falling exponentials τ_2_ = 12 ± 1.1 (11 ± 6.8) minutes. When the IED rate curves were averaged over mice (Figure 6C, black line) and fitted with the sum of two exponentials (Figure 6F, black line), the time constants were τ_1_ = 4.9 and τ_2_ = 12.2 minutes (Figure 6G).

Next, we analyzed 2-day periods before and after the seizures which displayed a > 2-day seizure-free period before or after them, respectively (Figure 6H). After the seizure, there was not only the previously mentioned fast transient increase (which appears as a sharp spike right after the seizure in the Figure 6H and Supplementary Figure 6) but also a much longer period of slightly increased IED rate 1 to 2 days after the seizure. Before the seizure, we observed a mild decreasing trend which, however, might be identical to the later stage of the after-seizure profile from the previous seizure (Figure 6I). We fitted the after-seizure data with a sum of three exponentials (increasing, decreasing, increasing) with τ_1_ = 1.8 ± 0.58 (0.96 ± 1.5), τ_2_ = 12 ± 6.9 (3.9 ± 6.6), and τ_3_ = 24 ± 7.2 (14 ± 25) hours (Figure 6K, L and Supplementary Figure 7). For the fit of the averaged IED curves over mice, it was with τ_1_ = 0.6, τ_2_ = 2.4, and τ_3_ = 22.3 hours (Figure 6L). The individual mice data can be found in Supplementary Figures 6 and 7.

### Simulation of IED rate

In the previous analysis, we identified a characteristic shape of the post-ictal IED rate temporal profile. Since we did not observe any characteristic changes in the IED rate ahead of seizures, we speculated that the long-term changes in the IED rate could be attributed to the time-dependent cumulative effect of the seizure-related increase in IED rate. To test this hypothesis, we simulated a long-term IED profile by causal filtering of the seizure rate data (see Methods and Figure 7A). We tested two approaches. In the first, subject-specific approach, we simulated the data using a filter derived only from the data of the given mouse (subject-specific approach, green in Figure 7). In the second approach, we used a filter derived from the averaged data over all mice, i.e. the black curve in Figure 6K (blue in Figure 7).

**Figure 7:**
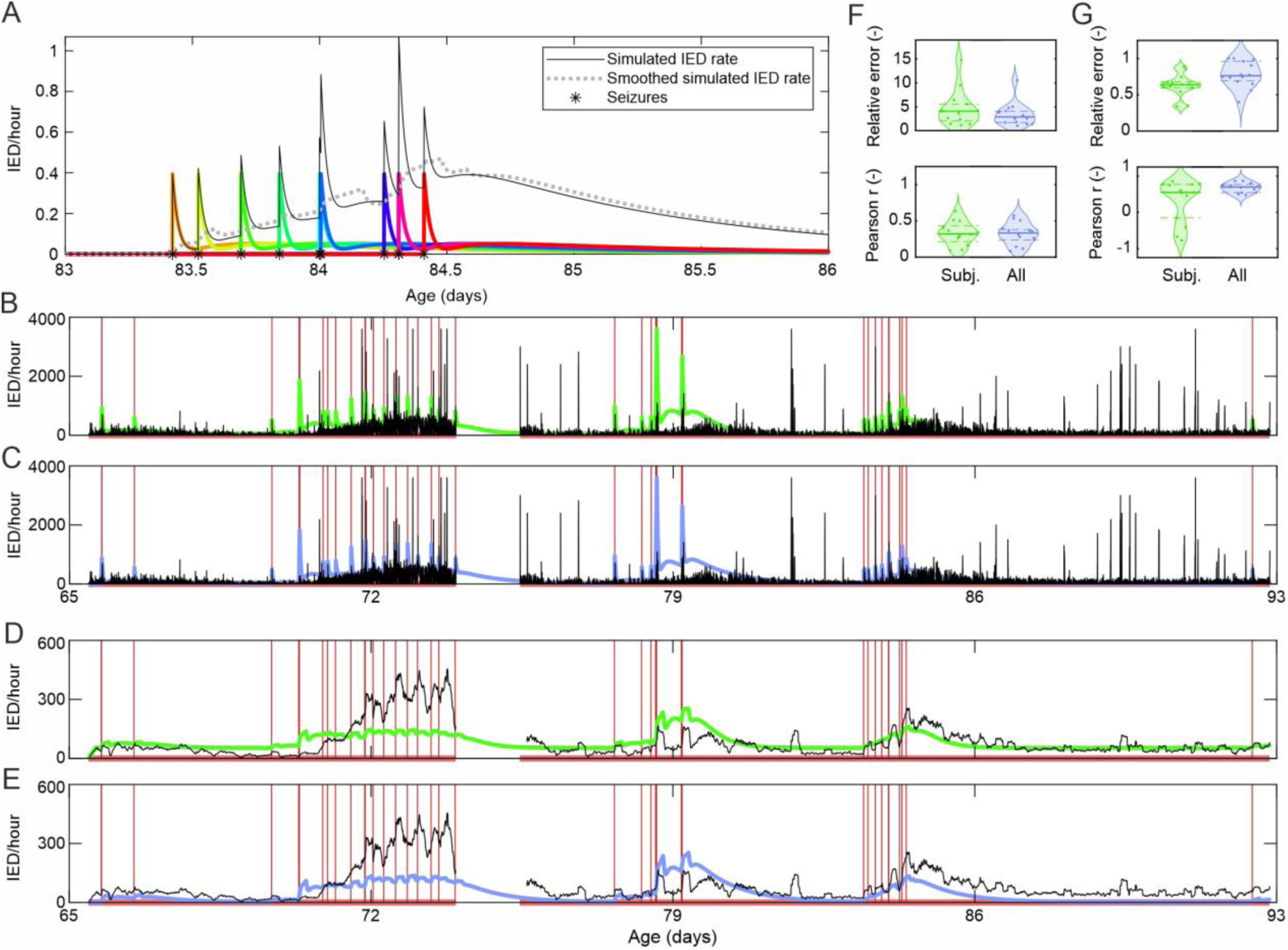
Simulation of the long-term profile of IED rate. (A) Example or a seizure cluster from Mouse 13 as an explanation of how each seizure (non-zero element in the seizure rate signal, marked by *), results in the presence of an impulse response (rainbow colors). The impulse responses add up to create a simulated IED rate (black curve). The dashed gray line is the black curve smoothed by 4-hour-long moving average filter. (B) Example of the original (black line) and simulated (green line) IED rate analyzed in 1-minute bins using the subject-specific impulse response. The red vertical lines represent seizures and the red line at the bottom indicates presence of the recording. Note a significant dropout on day 74. (C) Same as B but with the impulse response derived from the averaged IED rate over all mice. (D, E) Same as B, C but with 4-hour-long moving average smoothing of both original and simulated IED rate. (F) Relative error and Pearson’s correlation coefficient between the original IED rate and the simulated one using subject-specific impulse response derived from the mouse’s post-ictal IED rate (green) or impulse response derived from the mean post-ictal IED rate of all mice (blue). (G) Same as B but for smoothed versions of both original and simulated IED rate.

We also tested using either the raw filtered seizure rate (Figure 7B, C, F) or filtered seizure rate with additional smoothing by a causal 4-hour moving average (Figure 7D, E, G). This gave us four types of simulation. We evaluated the similarity of the simulated IED rate to the original IED rate by two metrics. The first one was Relative error, which was the root mean squared error normalized by the root mean squared value of the original IED rate. The second was Pearson’s correlation coefficient *r*. The four methods performed significantly differently in terms of both Relative error and Pearson’s *r* (p<0.001 for both, Kruskal-Wallis test). The smallest Relative error 0.62±0.04 (0.63±0.13) was observed for the smoothed data obtained with the subject-specific filter. The highest Pearson’s *r* = 0.68±0.04 (0.7±0.21) was observed for the smoothed data obtained with the population-derived filter. Thus, we conclude, that the smoothing emphasizes the trends which are common to the original and simulated IED rate. The subject-specific filter seems to perform similarly to the population-derived filter. On the smoothed data, the population-derived filter has a slightly higher Relative error but the Pearson’s *r* is better. Notably, in three mice, the subject-specific filter yielded negative Pearson’s *r* due to inadequate fitting of the noisy after-seizure IED rate.

## Discussion

This study provides novel information about the spontaneous seizure dynamics in the model of FCD-related epilepsy and, more importantly, about the nature of long-term and seizure-related dynamics of interictal epileptiform activity. The study demonstrates a two-peak profile of seizure-induced changes in IED activity that can explain long-term fluctuations in IED rate and their mutual relationship with seizures.

On the longest timescale, the whole recording of several weeks or months duration, we did not observe consistent increase or decrease in the seizure rate and seizure signal power with time but the seizure duration increased. One can argue that a longer monitoring period could potentially reveal a trend in seizure rate since Williams et al. (2009) reported that the increase in seizure rate in the kainate model of temporal lobe epilepsy (TLE) was not apparent until the second month of recording in some subjects. We recorded multiple subjects for almost two months and approximately half of them showed an increasing trend and the other half a decreasing trend in seizure rate, similarly to the mice with shorter recording (Supplementary Figure 1). This renders it rather unlikely that having two-month or longer recordings from all the mice would uncover a significant and consistent trend. Rats with pilocarpine-induced TLE showed a long-term decrease in IED rate possibly due to gradual cell loss (Behr et al., 2017). We observed an increase in IED rate over the whole recording period. Thus, the FCD-related epilepsy in our model was progressive in terms of seizure severity (duration) and interictal activity. Future studies should evaluate if the progression also accompanied by a gradual cognitive decline and changes in behavior.

Similarly to the vast majority of longitudinal studies on seizure occurrence in patients (Balish et al., 1991; Bauer and Burr, 2001; Griffiths and Fox, 1938; Sunderam et al., 2007; Tauboll et al., 1991) and animals models of epilepsy (Bajorat et al., 2011; B. L. Chang et al., 2018; Dudek and Staley, 2011; Eskikand et al., 2025; Goffin et al., 2007; Grabenstatter et al., 2005; Gregg et al., 2020; Hawkins and Mellanby, 1987; Kadam et al., 2010; Kudlacek et al., 2021; Mazzuferi et al., 2012), we observed seizure occurrence in clusters. Since the laboratory animals are kept in constant conditions, such studies, including ours, indicate that the seizure clustering is not driven by external factors, such as weekly cycle, but is generated endogenously. It is not known to what extent the seizure clustering is driven by physiological rhythms such as estrus cycle in females (Li et al., 2020) and to what extent by the epileptic processes (Kudlacek et al., 2021). At the level of individual clusters, we observed that seizure rate did not change but seizure duration, signal power and IED rate increased along the cluster which can be viewed as an increase in seizure severity. These observations are partly similar to our findings in the tetanus toxin model of TLE where clusters were characterized by an increase in spatial propagation of ictal activity and behavioral severity of the seizures over the course of a cluster (Kudlacek et al., 2021). In the tetanus toxin model of TLE, seizure durations were also previously observed to be linked to the seizure rate and consistently change along each seizure cluster (Eskikand et al., 2024). In agreement with clusters in human patients, we observed that the last seizure of a cluster was longer than the previous within-cluster seizures (Ferastraoaru et al., 2016). A possible mechanism behind the seizure clustering is that the shorter and less severe within-cluster seizures are insufficient to trigger postictal inhibitory processes. The longer or more generalized seizures, however, trigger these anti-seizure processes, which leads to the termination of the seizure cluster (Ferastraoaru et al., 2016; Kudlacek et al., 2021). Thus, the data from our FCD model support the view that the seizures themselves are implicated in the governance of the long-term dynamics of seizure occurrence and clustering. The knowledge of the seizure characteristics could, in theory, be leveraged for a clinically highly relevant task of prediction of seizure recurrence (Ferastraoaru et al., 2016).

We observed an increasing trend in IED rate over the course of the cluster, which can be in agreement with the human chronic iEEG data (Baud et al., 2018) and data from rats with TLE (Baud et al., 2019) and possibly rats with cortical lesions due to hypoxic-ischemic injury (Kadam et al., 2010). Baud et al. (2019, 2018) reported multidien cycles of IED rate not only in association with seizures but also during seizure-free periods. In contrast, our results do not show cyclic patterns of IED rate during seizure-free periods (Supplementary Figure 1). This difference could be attributed to different types of epilepsy (TLE rats vs. FCD mice). Another possible reason for this difference could, in theory, be that in the patients (Baud et al., 2018) and TLE rats (Baud et al., 2019), there could be subclinical events driving the IED rate cycles which did not meet the criteria to be classified as seizures in those studies.

The analysis of circadian rhythms revealed a higher seizure incidence in the afternoon and evening with some inter-individual variation and the mean at 4 p.m. under non-reversed light regime (switching on at 6 a.m. and off 6 p.m.). Since the mice are nocturnal animals, the seizures in our mice may correspond to awakening-related seizures observed in some patients with extra-temporal lobe epilepsy (Quigg et al., 1998). Because our mice had their FCD lesions most often in the frontal and parietal cortices, our data are in line with the findings of Durazzo et al. (2008) who observed the highest likelihood of frontal and parietal lobe seizures between 4 a.m. and 7a.m. Analysis of chronic ambulatory as well as shorter EEG recordings revealed a higher incidence of neocortical seizures during the night, whereas temporal lobe seizures peaked during the day (Hofstra et al., 2009; Mirzoev et al., 2012; Spencer et al., 2016). Our data are also in line with the results of Wang et al. (2022) who reported higher incidence of sleep-related epilepsy in humans with FCD compared to other etiologies. Our results also correspond to the data from a similar murine model of FCD where seizures displayed a higher seizure incidence during the light phase of the day (Almacellas Barbanoj et al., 2024). The translation between human and animal studies is, however, not straightforward. In humans, the temporal lobe and neocortical seizures have different preferential times of occurrence consistently across a large body of literature (Durazzo et al., 2008; Hofstra et al., 2009; Mirzoev et al., 2012; Quigg et al., 1998; Spencer et al., 2016; Wang et al., 2022). Meanwhile, in animal models, the seizures are more prevalent during the light period of the day regardless of the epileptic focus localization (Baud et al., 2019; Bertram and Cornett, 1994; Hellier and Dudek, 1999; Pitsch et al., 2017; Raedt et al., 2009 for TLE models and Almacellas Barbanoj et al., 2024 and present data for neocortical FCD-related epilepsy). In contrast to the human patient (Baud et al., 2018; Karoly et al., 2016) and animal studies (Baud et al., 2019), we observed only a mild circadian modulation of IED rate in individual mice with a high inter-individual variability in the peak phase.

Next, we analyzed the changes before and after individual seizures. The most striking finding was a period of increased IED rate after seizures. Our observation is in agreement with data from TLE patients (Gotman and Marciani, 1985) and the tetanus toxin model of TLE (Hawkins and Mellanby, 1987), but in contrast to the pilocarpine and kainate model of TLE, where no difference between the pre-ictal and post-ictal IED rates was reported (Baud et al., 2019). In contrast to Karoly et al. (2016) and Levesque et al. (2011), we did not observe any change in IED rate preceding the seizures. Our results are also in a seeming contrast with the human chronic iEEG studies, where seizures were shown to occur mostly on the rising phases of the IED rate fluctuation, implying that the IED rate was increasing already before the seizure (Baud et al., 2018; Leguia et al., 2021; Maturana et al., 2020). In these studies, however, non-causal filtering or cycle phase estimation method was employed. Thus, those algorithms use the information from the future and possibly may project the post-seizure IED rate increase to the pre-seizure period.

The observation of early and late kinetics in seizure-related two-peak IED rate profile after the seizures is particularly intriguing. Our modeling has shown that cluster-associated changes in IED rate can be reproduced using only seizure timing and the two-peak IED rate response. Notably, the slow component appears crucial in producing a cumulative temporal effect that contributes to the multi-day dynamics observed during seizure clusters. From a pathophysiological perspective, the fast and slow IED changes likely reflect distinct underlying cellular and network mechanisms regulating excitability. Postictal activation of epileptiform discharges is commonly observed in humans (Kaibara and Blume, 1988; King et al., 1998; So and Blume, 2010). Immediately following the seizure termination (Lado and Moshé, 2008), seizure-generating regions often exhibit profound disruption in extracellular space homeostasis and metabolic imbalance (Blauwblomme et al., 2014; Lux and Heinemann, 1978; Raimondo et al., 2015), including persistently elevated extracellular potassium levels or lowered calcium, conditions that may promote IEDs specifically in those regions that generated ictal activity actively (Kaibara and Blume, 1988). The delayed increase in IED rate, typically occurring about one day post-seizure, may involve additional processes that elevate network excitability and potentially promote a state of heightened seizure risk. This secondary rise in IED activity may be attributed to slow-developing pathological changes, such as a progressive weakening of anti-seizure mechanisms (Lado and Moshé, 2008) or the emergence of seizure-induced pro-excitatory processes, including neuroinflammation, blood-brain barrier disruption, or glial dysfunction (Vezzani et al., 2011; Vezzani and Granata, 2005). Seizures are known to trigger the release of pro-inflammatory cytokines, like IL-1β, IL-6, and TNF-α, with a latency of several hours (De Simoni et al., 2000; Vezzani et al., 1999). This can be associated with microglial activation and astrocyte dysfunction. Conversely, anti-inflammatory responses are also induced but generally exhibit a more delayed onset. This temporal mismatch may play a key role in shaping delayed postictal IED activation, cluster formation and cluster-related IED profile (Jiruska et al., 2023; Vezzani et al., 2011).

To describe quantitatively the post-seizure IED rate, we fitted the 2-hour post-seizure IED rate curve with a sum of two exponentials, faster increasing and slower decreasing. A 2-day post-seizure IED rate curve could be fitted with a sum of three exponentials. We then simulated IED rate fluctuations using the three-exponential model and seizure timings. We discovered that when both original and simulated IED rate are smoothed by a 4-hour filter, the simulated data copy the fluctuations of the original IED rate. The correlation of the simulated and original IED rate was even better when an average IED rate response over all mice was used instead of individual mice data to derive the three-exponential model. These results suggest that for seizure forecasting or prediction, the IED rate may carry low additional information to the information contained in seizure records, provided the three-exponential model is known. Thus, the IEDs may be a mere reflection of seizure dynamics which would explain why some studies used IED rate for seizure forecasting (Maturana et al., 2020; Proix et al., 2021) whereas another study could forecast future seizure risk using seizure records only (Gleichgerrcht et al., 2022; Karoly et al., 2020).

Future studies should make use of multichannel data and investigate the epileptic networks since the connectivity patterns were reported to hold good promise for seizure forecasting (Khambhati et al., 2024; Lehnertz et al., 2023; Schroeder et al., 2023). We recorded at a low sampling rate of 2000 Hz with anti-aliasing filter cutoff at 500 Hz which prevented analysis of fast ripples which have the frequency of 600 Hz in this model of epilepsy (Chvojka et al., 2024). Another set of features which would be worth investigating are the markers of critical slowing down such as signal variance, lag-1 autocorrelation, skewness and spatial correlation (W. C. Chang et al., 2018; Maturana et al., 2020).

### Conclusion

This study demonstrated that seizures in a realistic mouse model of FCD-related epilepsy follow principles similar to those observed in other epilepsy models, including their progressive nature, temporal clustering, and modulation by circadian rhythms. We found that changes in seizure features are often mirrored by corresponding temporal fluctuations in the rate of IEDs. Specifically, seizure clusters are associated with a sustained increase in IED rate, which gradually returns to baseline after the cluster resolves. Notably, the rise in IED rate does not precede the onset of the cluster but rather follows its initiation. These changes in IED activity appear to be driven by individual seizure-related increases, exhibiting both fast and slow kinetic components. In particular, the processes responsible for the slow rise in IED rate may reflect epileptic network alterations that accumulate with repeated seizures, contributing to multi-day dynamics in IED rate and cluster formation. Future studies focusing on the cellular and network mechanisms underlying the slow postictal seizure-associated IED increases may be key to identifying preventive therapeutic strategies to avert cluster formation. The multiscale properties of network dynamics derived from seizures and seizure-related IED changes still hold valuable insights for the development of seizure forecasting approaches and improved monitoring of disease activity.

## Author contribution

Conceptualization: J.K., P.J.; Methodology: J.K., M.B., P.J.; Resources: J.O., M.B.; Investigation: J.C., T.R.; Software: J.K., J.C.; Data curation: J.K., J.C., T.R.; Formal analysis: J.K.; Validation: M.K., S.K.; Visualization: J.K., J.C.; Supervision: J.O., M.B., P.J.; Funding acquisition: J.K., O.N., P.J.; Writing – original draft: J.K., P.J.; Writing – review & editing: J.K., J.C., M.K., S.K., T.R., O.N., J.O., M.B., P.J.

## Data availability

Recorded data and analytical tools used in this study are available from the corresponding author upon request.

