## Supplementary figures for "Interictal activity fluctuations follow rather than precede seizures on multiple time scales in a mouse model of focal cortical dysplasia"

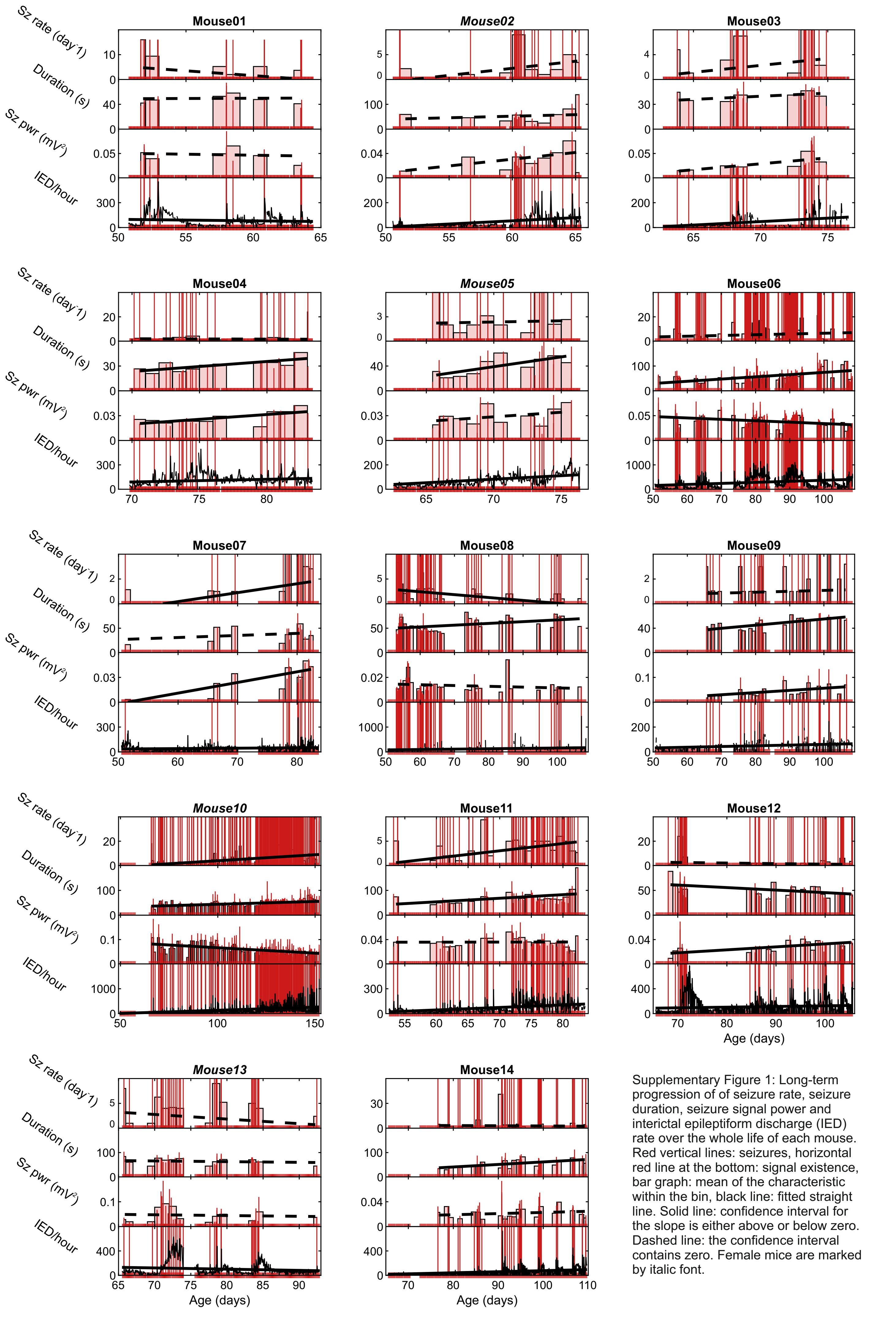


Supplementary Figure 1: Long-term progression of of seizure rate, seizure duration, seizure signal power and interictal epileptiform discharge (IED) rate over the whole life of each mouse. Red vertical lines: seizures, horizontal red line at the bottom: signal existence, bar graph: mean of the characteristic within the bin, black line: fitted straight line. Solid line: confidence interval for the slope is either above or below zero. Dashed line: the confidence interval contains zero. Female mice are marked by italic font.

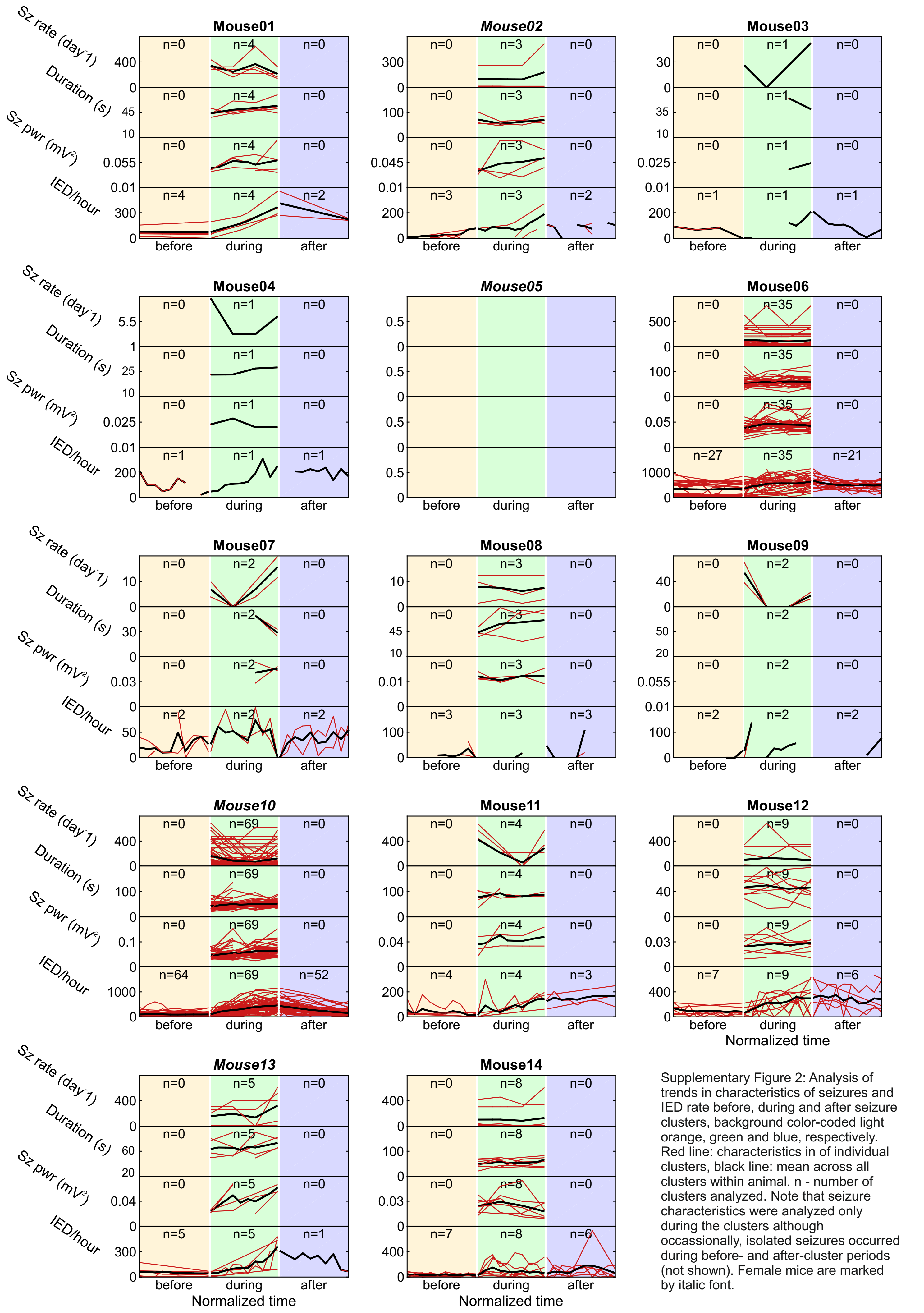


Supplementary Figure 2: Analysis of trends in characteristics of seizures and IED rate before, during and after seizure clusters, background color-coded light orange, green and blue, respectively. Red line: characteristics in of individual clusters, black line: mean across all clusters within animal. n - number of clusters analyzed. Note that seizure characteristics were analyzed only during the clusters although occassionally, isolated seizures occurred during before- and after-cluster periods (not shown). Female mice are marked by italic font.

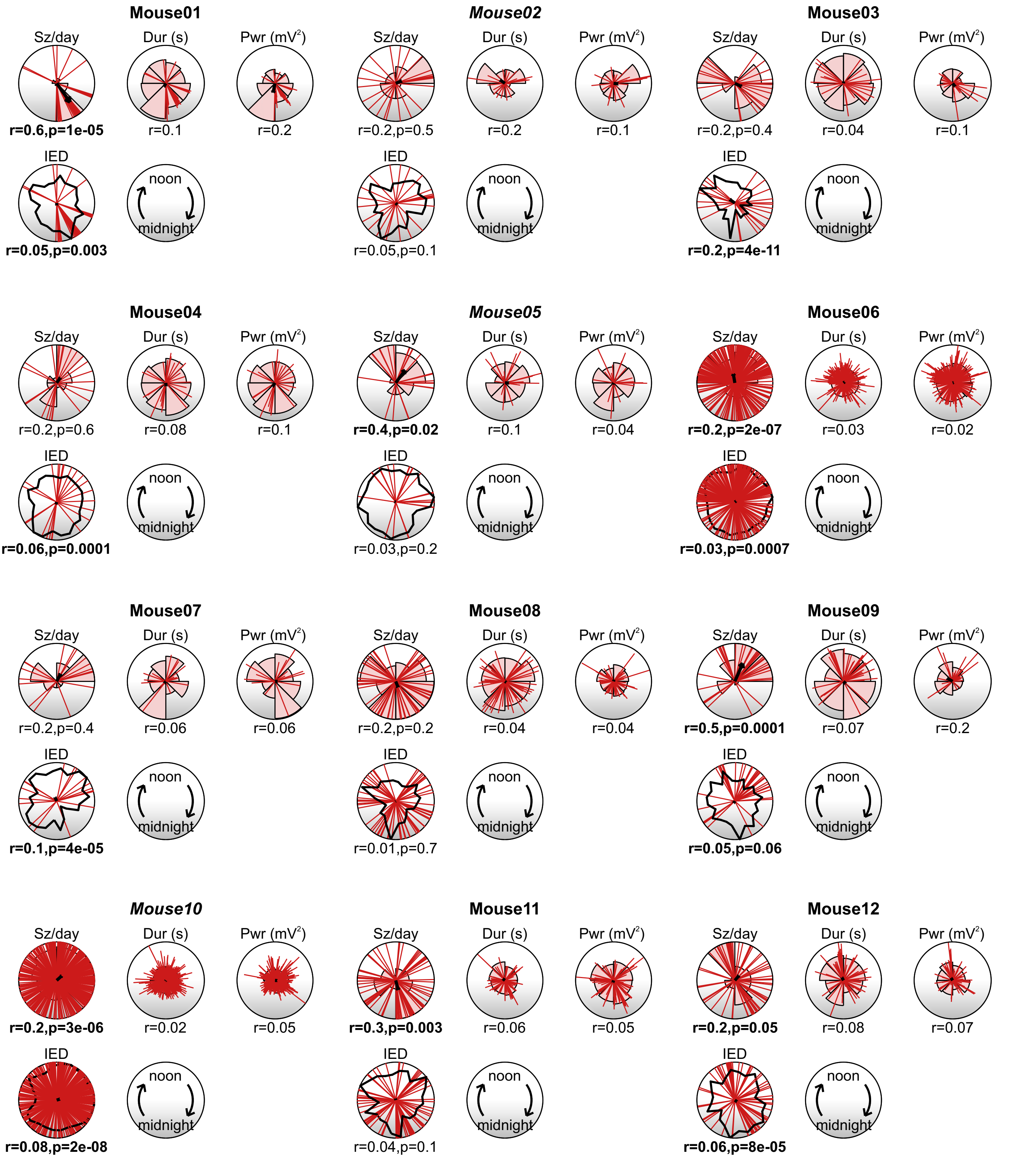

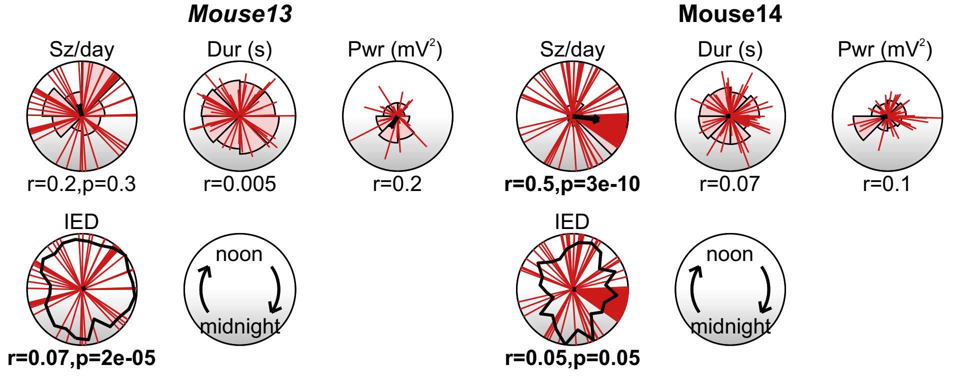


Supplementary Figure 3: Circadian variation of seizure characteristics and IED rate in individual animals. Red lines: seizures. Circular bar graphs: mean of the characteristic within the bin. Black arrow: mean resultant vector. For IED rate the distribution is shown by a black curve with the seizures underneath. Midnight is at the bottom, noon at the top and time goes clockwise. r: length of the mean resultant vector. p: p-value of Rayleigh test. Bold font: $p < 0.05$. Note that for the Rayleigh test was performed only for seizure and IED occurrence data and not for seizure characteristics. Female mice are marked by italic font.

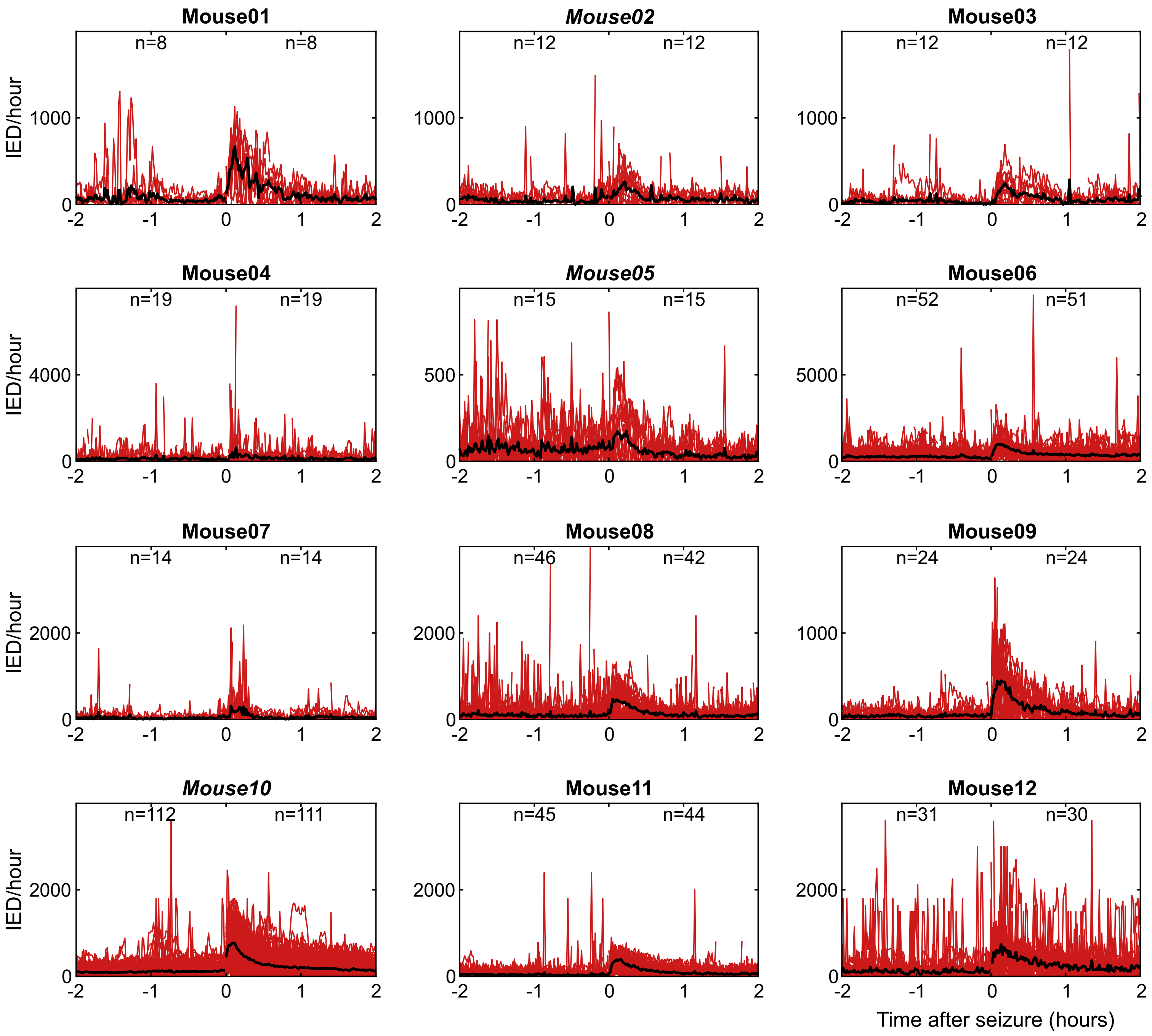


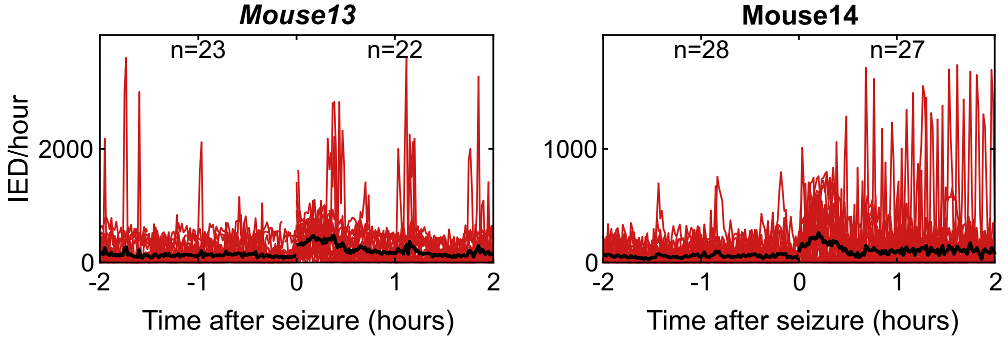


Supplementary Figure 4: IED rate two hours before and after seizure, evaluated in 1-minute bins. Red curves: Individual seizures. Black curve: Mean over seizures. characteristics. Female mice are marked by italic font.

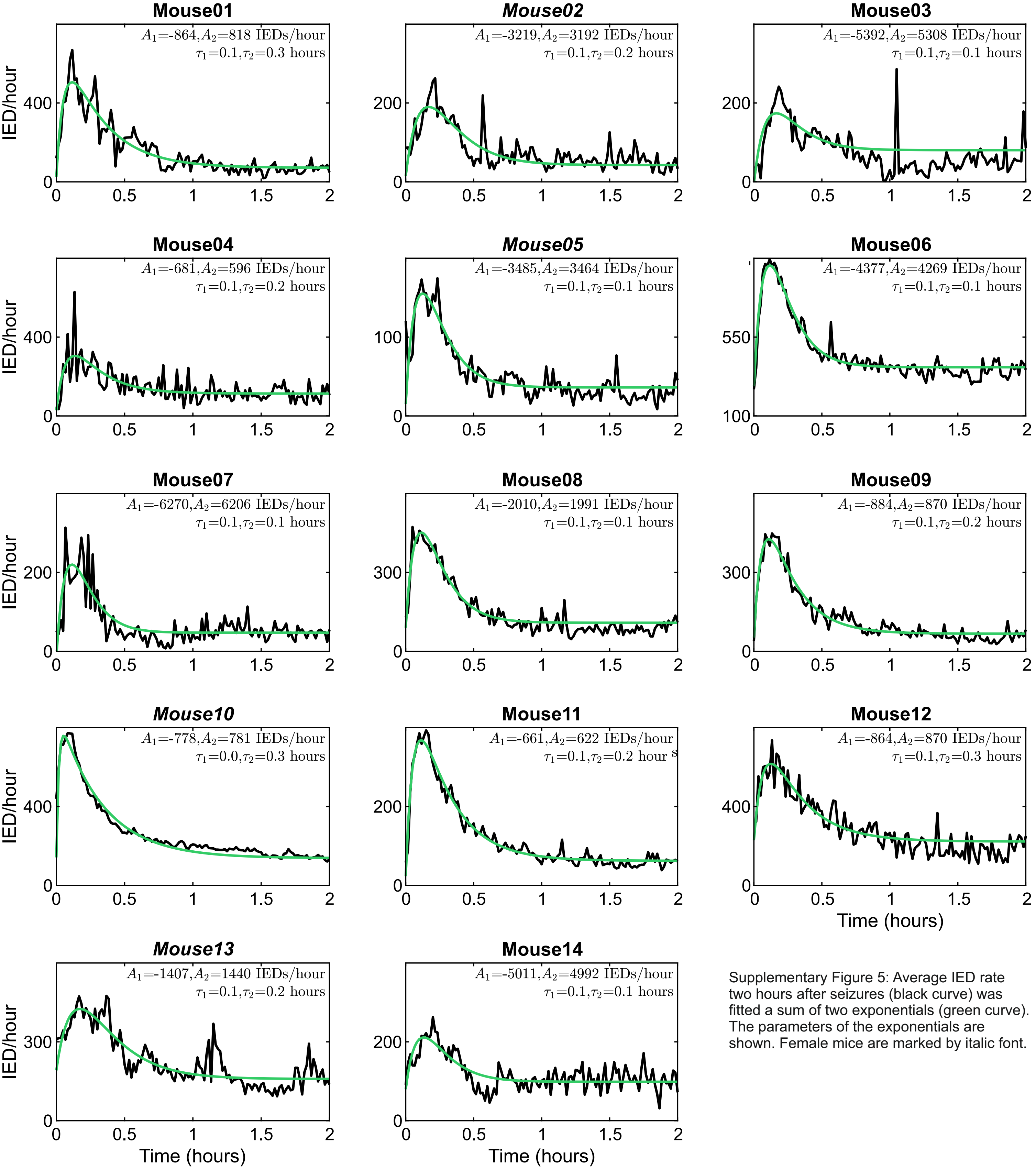


Supplementary Figure 5: Average IED rate two hours after seizures (black curve) was fitted a sum of two exponentials (green curve). The parameters of the exponentials are shown. Female mice are marked by italic font.

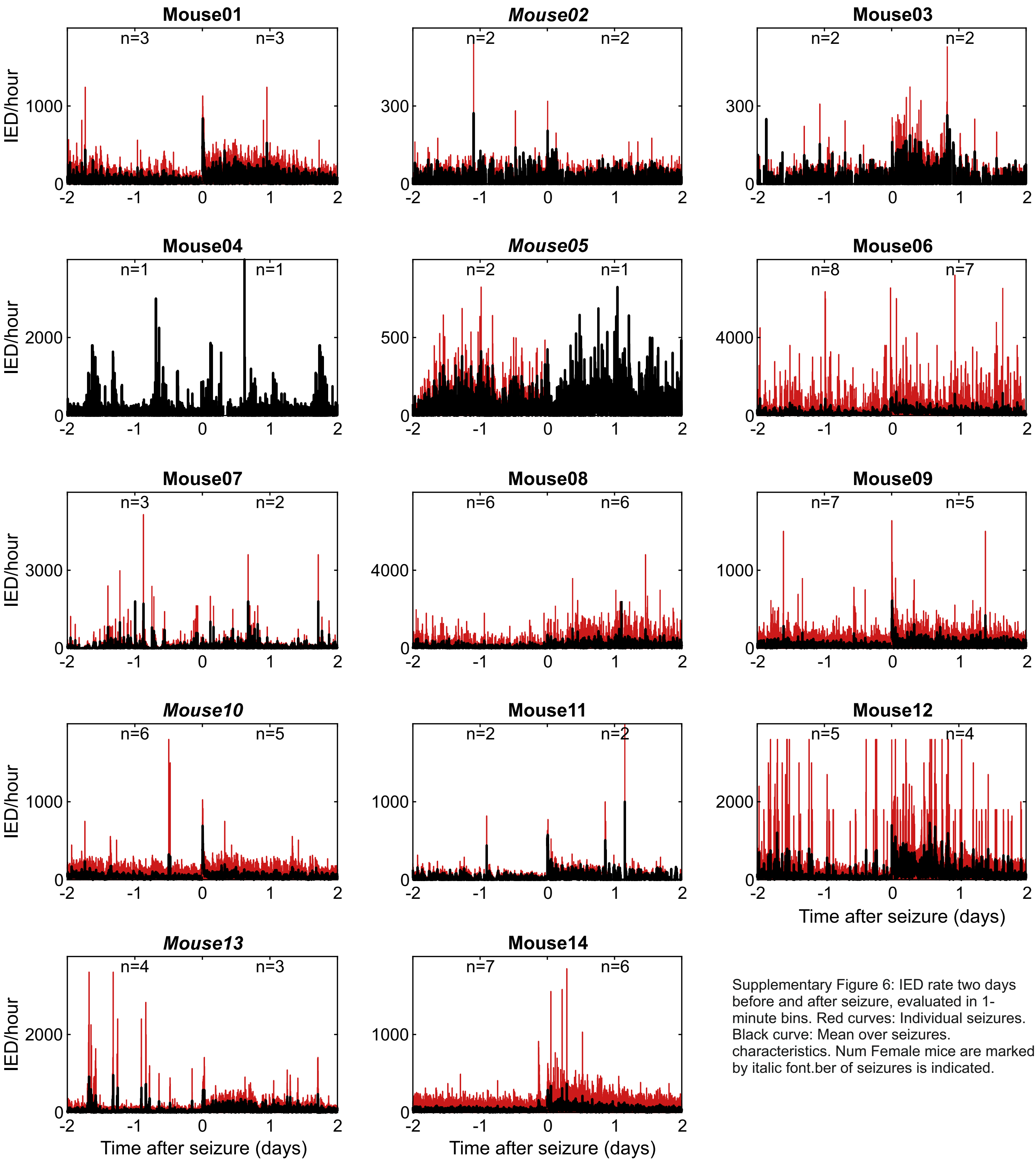


Supplementary Figure 6: IED rate two days before and after seizure, evaluated in 1-minute bins. Red curves: Individual seizures. Black curve: Mean over seizures. characteristics. Num Female mice are marked by italic font.ber of seizures is indicated.

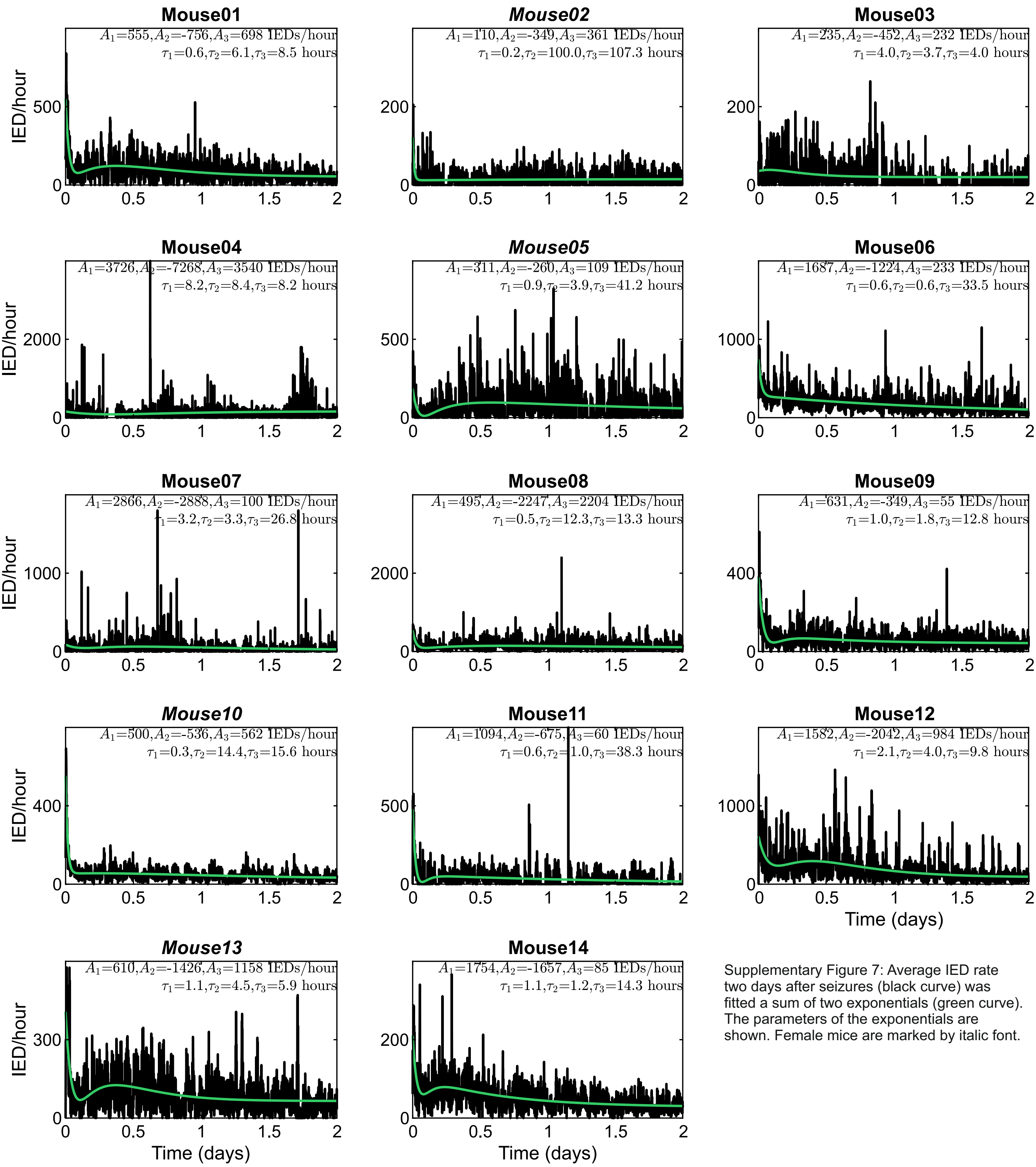


Supplementary Figure 7: Average IED rate two days after seizures (black curve) was fitted a sum of two exponentials (green curve). The parameters of the exponentials are shown. Female mice are marked by italic font.

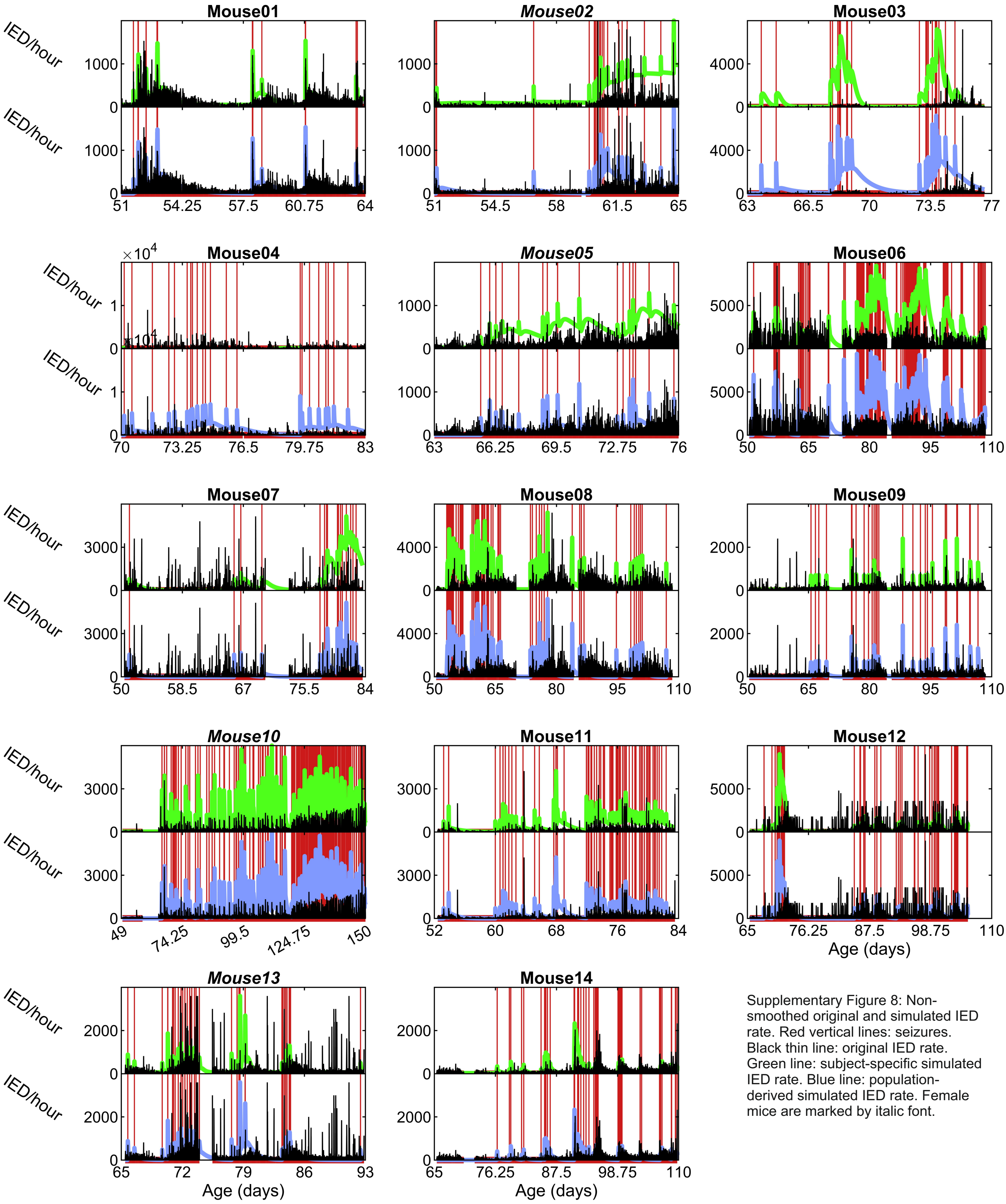


Supplementary Figure 8: Non-smoothed original and simulated IED rate. Red vertical lines: seizures. Black thin line: original IED rate. Green line: subject-specific simulated IED rate. Blue line: population-derived simulated IED rate. Female mice are marked by italic font.

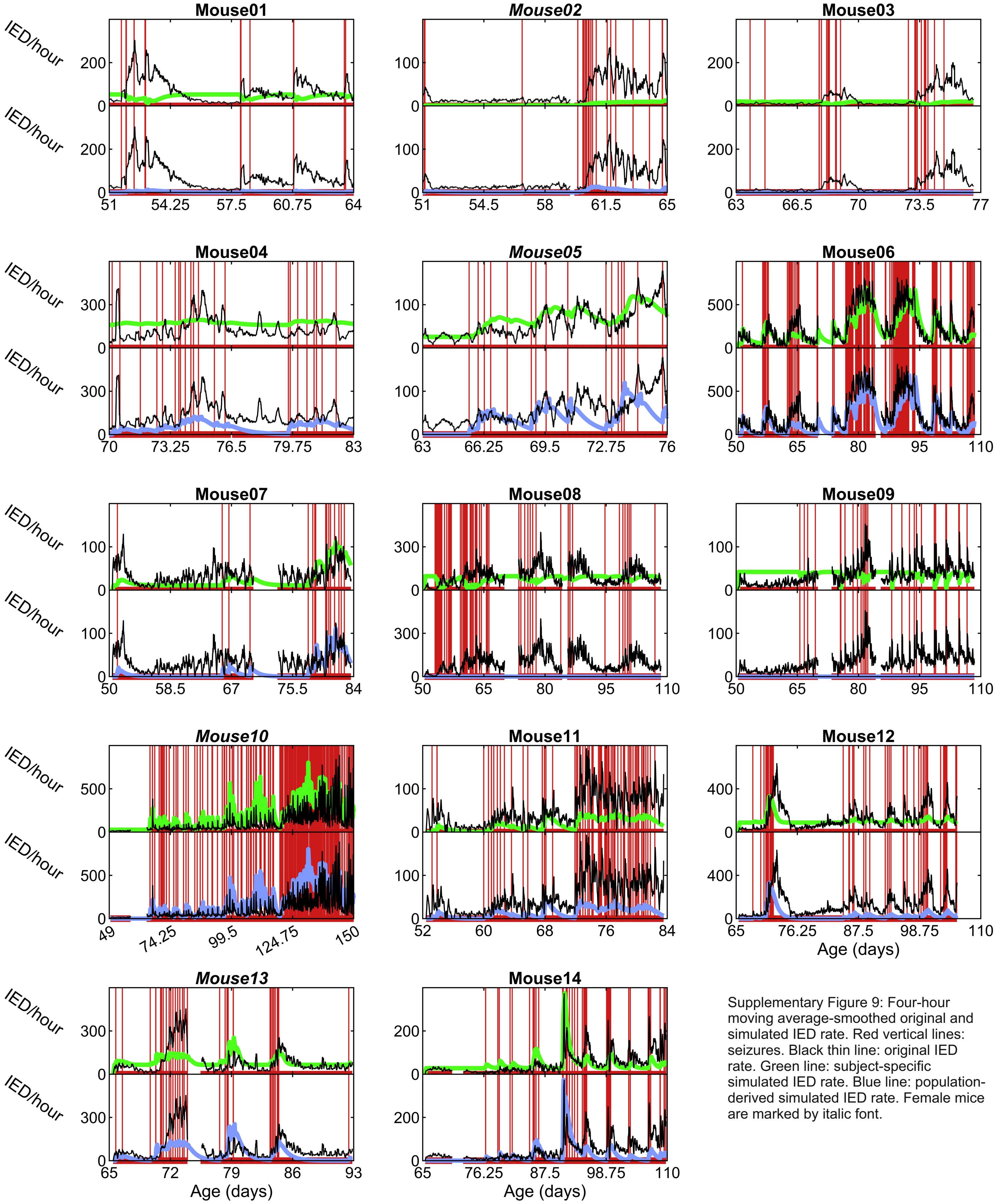


Supplementary Figure 9: Four-hour moving average-smoothed original and simulated IED rate. Red vertical lines: seizures. Black thin line: original IED rate. Green line: subject-specific simulated IED rate. Blue line: population-derived simulated IED rate. Female mice are marked by italic font.
